# Somatic copy number mutations contribute to fitness in transplantation models of spontaneous human breast cancer metastasis

**DOI:** 10.1101/2025.10.27.684911

**Authors:** Hoa Tran, Gurdeep Singh, Hakwoo Lee, Damian Yap, Eric Lee, William Daniels, Farhia Kabeer, Ciara H O’Flanagan, Vinci Au, Michael Van Vliet, Daniel Lai, Elena Zaikova, Sean Beatty, Esther Kong, Shuyu Fan, Jessica Chan, Hoang Quan Dang, Viviana Cerda, Teresa Ruiz de Algaza, Andrew Roth, Samuel Aparicio

## Abstract

The contribution of somatic gene dosage mutations (CNA) to breast cancer metastasis remains poorly defined. Using 9 transplantable human neoadjuvant-naive triple-negative breast cancer xenografts, we studied the fitness of copy number clones in spontaneous metastasis from orthotopic transplant sites. Metastatic site preference was strongly patient-dependent, and the emergence of metastases exhibited a general trend toward slower growth at the orthotopic site. In our models, single-cell whole-genome sequencing of primary and metastatic sites showed that distant metastases were most often the result of minor prevalence clones at the orthotopic site, suggesting that some metastatic phenotypes may be weakly negatively fit at the primary site. We validated the existence of a fitness hierarchy of copy number clones using a previously established paradigm of remixing and retransplanting clones. Single-cell clone analysis of competitive repopulation and re-emergence of metastases showed that CNAs arising in cancer evolution can mediate metastatic fitness. Moreover, some clones displaying strong metastatic tendency exhibited weaker survival at the primary site, consistent with the notion that metastatic phenotypes could have a fitness cost at the primary site. Finally, we conducted RNA-seq analysis combined with DriverNet analysis to dissect the contribution of CNA-mediated versus genome-independent transcriptional states. CNA mutations appeared to contribute strongly to transcriptional differences between clones. Among clones of high metastatic potential, we observed CNA-mediated and CNA-independent convergence on pathways such as epithelial-mesenchymal transition (EMT), established as mediators of metastatic cell survival at distant sites. Taken together, our data point to a contribution of CNA-mediated cancer evolution to metastatic states and identify distant-site context as a key determinant of CNA-mediated fitness.

## INTRODUCTION

Metastatic progression in breast cancer remains an incurable stage of disease, associated with therapeutic resistance and evolutionary plasticity. Breast cancers are genomically unstable with mutations ranging from base substitutions to whole-chromosome aneuploidy. Previous studies have demonstrated the dual contribution of single-nucleotide mutations (SNV) and large-scale copy number alterations (CNA), including whole-genome duplication (WGD)^1,2^ in tumour evolution The role of mutations in the emergence of primary breast cancer has been extensively documented^3–5^; however, the contribution of chromosomal-scale gene dosage mutations and genome doubling to the breast cancer metastatic states has only been recently explored^6,7^. Recent studies have shown that the metastatic states in breast cancer inherit phenotypes established early during breast cancer ontogeny, but also that ongoing CNA mutational processes, including WGD, appear to increase the genomic complexity of metastatic disease. The evolutionary transition of primary site tumour cells to site-specific cancer fitness is driven by both genetic and non-genetic processes ^8^, which allows metastatic cells to attain differential transcriptional properties fitter for migration, invasion, and colonization of other organs of the body^9–11^. Tumour heterogeneity plays a crucial role in the ontogeny of a mixed population with differential clonal fitness landscapes and differential properties, including drug-resistance^12^.

Triple-negative breast cancer (TNBC) is the most clinically and genomically heterogeneous breast cancer subtype. PDX models of breast cancer have been shown to capture key intrinsic properties^13–15^, especially for TNBC, where engraftment is a negative prognostic factor^16^. To measure the fitness characteristics of metastatic clones, we investigated the role of gene dosage mutations and transcriptional forces in TNBC patient-derived xenografts (PDX) models of metastasis. These orthotopic PDX models were established from neoadjuvant-naive primary breast tumours from 9 independent donors (Pt.1-9) and propagated via serial transplantation in mice. Following primary tumour resection (PTR) of the orthotopic PDX, capable models produced spontaneous metastases in vivo without experimental re-injection, demonstrating an intact intrinsic metastatic programme. PDX models such as these have been shown to recapitulate the genomic and phenotypic heterogeneity of human patients^17,18^, and have been validated as useful models of metastasis dynamics in human breast cancers^19^. We have recently shown that drug resistance to common breast cancer chemotherapeutic agents, such as platinum, occurs in part due to clonal selection of gene dosage mutations^20^ and in part through clone-independent transcriptional programs^21^. Here, we adopt an analogous approach, using single-cell genome sequencing of spontaneous metastases of PDXs, to identify the relative contribution of gene dosage mutations to metastatic fitness *in vivo*.

## RESULTS

### Spontaneous metastasis patterns from human orthotopic primary site transplants are patient-specific and inversely related to primary growth rate

We investigated 9 neoadjuvant-naive TNBC patient biopsy samples with known engraftment capacity^(2, 16, 18)^ in immunodeficient mice to assess the spontaneous metastatic capacity after resection of the orthotopic primary tumour without experimental re-injection. As expected, all 9 lines developed orthotopic site tumours in multiple independent transplants (orthotopic primary PDX tumours) and maintained hormone receptor status (Supplementary Table 2, Supplementary Figures 1 and 2). Tumours were allowed to grow to the maximum allowed tumour volume of 1000 mm³, after which they were surgically removed (primary tumour resection, Figure 1A). Mice were then monitored for up to 50 weeks for metastases and orthotopic site regrowth, which we established through clinical signs of metastasis, palpation or PET-CT scans (39 scans over 9 lines) (Supplementary Table 1), and autopsy if no signs were observed after 50 weeks. Metastasis was detected in 5 patient PDX lines (Supplementary Table 1).

**Figure 1:**
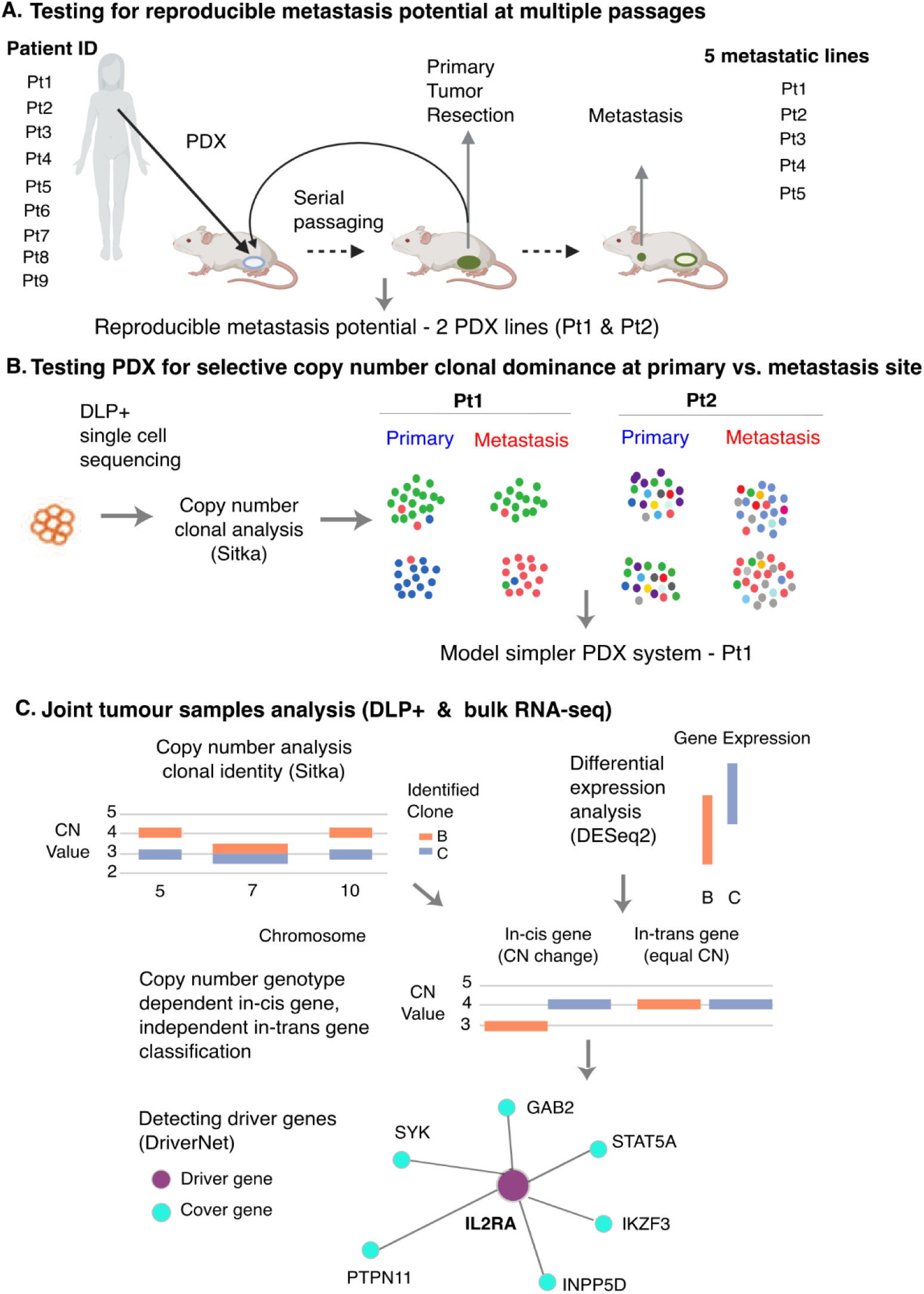
PDX study model to identify a highly competitive clonal system for individual decoding of genetic and non-genetic components. A) Experimental design to test the metastatic potential of neoadjuvant-naive primary breast cancer cells from patients Pt1-9 in immunodeficient mice over multiple passages (figure created using BioRender.com). PDX lines Pt1-5 developed metastasis. Of these 5, 2 PDX lines (Pt1 and Pt2) were selected for extended analyses based on reproducible metastatic potential. B) Cell profiles were captured using scWGS (DLP+). Copy number clonal analyses were performed using the Sitka package. Cell copy number profiles were used to identify the clonal identities and relative composition in both primary and metastasis samples. Each dot represents a cell with colours denoting the clonal identity - cell phenotype. Pt1 PDX transplantation model is selected for further experiments and downstream analysis based on its metastatic potential linked to its clonal evolution. C) Joint tumour populations were captured by DLP+ to measure copy number clonal profiles and bulk RNA-seq to identify differentially expressed genes. Genes were classified into copy-number genotype dependent in-cis genes or copy number genotype independent in-trans genes based on the overlap between their genomic regions and regions with copy number change in the DLP+ data. Driver genes were identified using the DriverNet package.

The pattern of metastasis appeared highly patient line specific (Supplementary Table 1), as has been previously documented for breast PDX tumours^13,22–24^, for example Pt1-SA919 only exhibited paraspinal site metastases in multiple replicate transplants (n=6) (Figure 2B), with a strong bias for upper thoracic/cervical region, whereas Pt2-SA535 exhibited lymph node, lung and liver metastasis, reproducibly (n=7) (Figure 2B). Metastatic potential varied across passages with no consistent trend: Pt1-SA919 only exhibited metastasis after passage X3 despite primary orthotopic site growth, Pt2-SA535 had metastatic potential in all passages, and Pt3-SA1142 lost metastatic potential and primary orthotopic site growth after passage X2. A key question concerns the relationship between organotypic metastasis in the PDX system and in corresponding TNBC patients. We observed that the development of metastasis in PDX was independent of patients’ metastasis status in 4 lines (Supplementary Table 1). Pt2-SA535 and Pt3-SA1142 developed metastases in PDXs, whereas patients remained disease-free up to 149 months (range 72-149 months) (Supplementary Table 1). On the other hand, the originating patient tumours of Pt6-SA604 and Pt7-SA501 metastasized in the patients but not in PDXs. Three PDX lines showed metastasis development in both PDXs and patients with different organ involvement (Pt1-SA919, Pt4-SA605, Pt5-SA609). Two lines showed no distant metastasis in both PDXs and patients (Pt8-SA1139, Pt9-SA1146). This divergence suggests that metastatic competence in PDXs arises from tumour cell-intrinsic properties, which may be further modulated by host microenvironmental factors. Contrasting theories suggest that rapidly dividing tumours may be more prone to metastasis^25^, whereas it has also been emphasized that expression of metastatic potential is a time-dependent phenomenon^26^. To test whether primary-site growth dynamics could predict metastatic potential, we next examined the relationship between primary orthotopic site growth rate and metastatic potential (Supplementary Table 3). We fitted a linear mixed-effects model with the cubic root of tumour volume as the response variable, the metastatic status as the fixed effect variable and treating each tumour as a random effect (see Methods). Across all lines, PDX transplants developing metastases exhibited a significantly slower orthotopic primary site growth rate (-0.075 Δ, p-value < 0.01, see Methods) compared with primary orthotopic site transplants with no metastatic development (Figure 2A-B). However, some interpatient heterogeneity was noted. For Pt1-SA919, the growth rate for orthotopic primary site transplant tumours that metastasized was slightly faster (0.056 Δ growth rate parameter with p-value = 0.007, see Methods) compared with Pt1-SA919 primary transplants that didn’t metastasize. Among PDX transplants assayed for mitotic cell fraction (Pt1-SA919, Pt2-SA535, Pt3-SA1142; Ki67, see Methods), PDX transplants that eventually developed metastases (29.5% ± 8.4% of Ki67+) showed a slightly lower fraction of proliferating cells than those that did not metastasize (32.1 ± 10.1% of Ki67+). However, the trend did not achieve statistical significance (KS test: D=0.16, p-value=0.2, see Methods)(Supplementary Figure 2D, Supplementary Table 2). These findings suggest that metastatic competence in our PDX models is not simply a function of rapid proliferation but instead reflects a fitness balance between local expansion and metastatic dissemination capacity.

**Figure 2:**
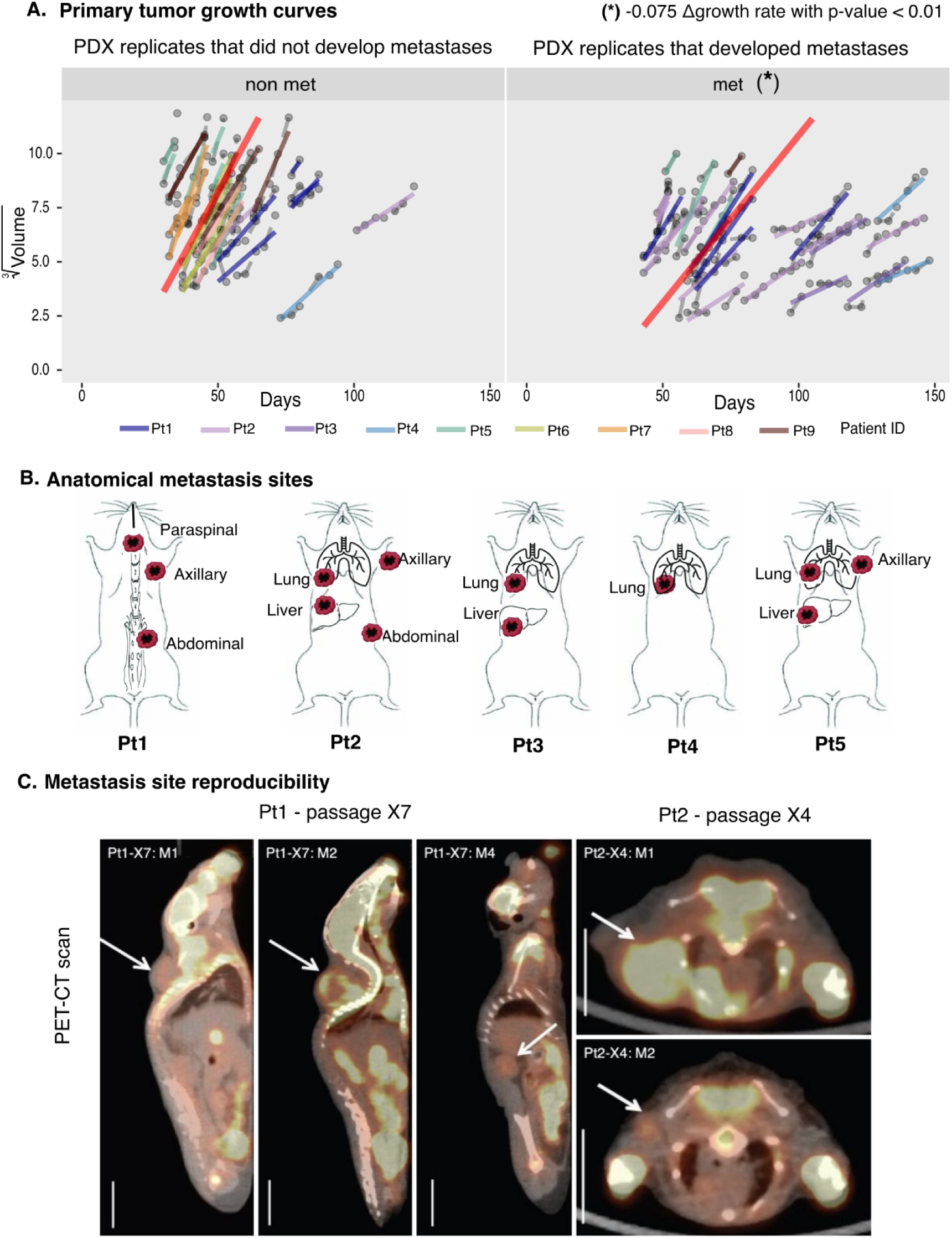
Replication rates and anatomical sites of metastasis. A) Relationship between primary site tumour growth rate and metastatic potential. Tumour volume against time post-implant (days) are shown for non-metastatic implants (left) and metastatic implants (right). Each implant is summarized with a fitted linear model shown in growth lines colored by patient ID. The omnibus linear model is shown in red. B) Schematic anatomical sites of reproducible metastasis over 5 PDX lines that demonstrated high metastasis reproducibility: Pt1-SA919, Pt2-SA535, Pt3-SA1142, Pt4-SA605, Pt5-SA609. C) Representative PET-CT images for PDX lines: Pt1-SA919 and Pt2-SA535 showing anatomical sites where metastases developed. PET-CT scan views of Pt1-SA919 passage 7 mouse M1, mouse M2, and mouse M4. Paraspinal masses at supraspinal and ventral spinal regions are identified (arrowheads). In Pt2-SA535 passage 4, a large mass was seen in the left axillary area of mouse M1, and in M2, a small mass uptake in the left axillary area and a large mass at the transplant site (arrowheads). Scale: 10 mm.

Finally, we assessed the reproducibility of metastatic potential after primary site resection in 3 to 4 different transplant replicates for each PDX passage. We observed that two patient lines, Pt1-SA919 and Pt2-SA535, resulted in frequent (≥50%) metastasis in at least 2 different passages, with reproducible, site-specific colonization patterns detected at stereotyped distant sites by PET/CT imaging scan (Figure 2C and Supplementary Table 1). Interestingly, we found that metastatic potential in Pt1-SA919 increased with passage number from 0% of primary implants in passage X3, to 50% in passage X4, and 100% in passage X7 (Supplementary Table 1), suggesting evolution of metastasis through intra-tumour heterogeneity clonal evolution. In contrast, Pt2-SA535 showed 100% metastasis development after primary site resection at all 4 primary passages from X4 to X7 (except X5, where only 3 / 4 primary tumours grew), suggesting metastatic potential was already established in Pt2-SA535 by passage X4, requiring no further evolution (Supplementary Table1). These results exemplify two evolutionary trajectories of metastatic competence within TNBC PDXs: progressive acquisition through intra-tumoural clonal evolution (Pt1-SA919) and constitutive maintenance of an established metastatic-competent clonal population (Pt2-SA535).

### Evolution of CNA-associated and CNA-independent metastatic potential identified from temporal single-cell genome sequencing

A key feature of human TNBC tumours is the dominant influence of gene dosage mutations (CNAs), which generate clonal complexity and pattern transcriptional responses. To test whether the large-scale genomic structural changes associated with CNA mutations drive the evolution of metastatic potential, we conducted single-cell whole-genome sequencing of orthotopic primary sites and metastatic deposits. We performed DLP+ single-cell whole-genome sequencing^27^ for the two patient lines, Pt1-SA919 and Pt2-SA535, exhibiting the most reproducible spontaneous metastatic patterns after primary tumour resection.

We first analyzed Pt2-SA535 passage X4 primary transplant replicates and their metastatic samples, assessing clonal composition and transplant replicates. Pt2-SA535 is a highly genomically unstable TNBC exhibiting hallmarks typical of this aggressive subgroup^2,20,21,28^. All primary passages of Pt2-SA535 (Supplementary Table 1) displayed metastatic potential with a mean interval ± sd to metastasis of 9.73 ± 3.26 weeks. We thus focused on passage X4, comprising 4 mouse replicates that showed the most reproducible metastasis sites, to ask whether there was evidence of clonal dominance associated with metastatic potential. We classified 4,407 high-quality single cell genomes, with a median ± sd of 304 ± 245 single cells per sample (Figure 3A, Supplementary Table 4) from 4 primary and 6 metastatic tumours (originating from axillary, inguinal, left and right axillary) using our previously described phylogenetic and tree cutting methods Sitka^20^(see Methods). Clonal copy-number heatmap of Pt2-SA535 (Figure 3A) and phylogenetic analysis (Figure 3B) revealed the clonal hierarchy of 11-15 CNA-defined clones organized into two main evolutionary branches, exhibiting primary tumour polyclonality. Metastatic tumours from replicate mice exhibited a lower degree of polyclonality with 1-5 major clones (cell proportions >= 10%) per metastasis (Figure 3C-D, Supplementary Table 5), with the majority of clones did not exceed 75% of the total cells, confirming that the primary sites were polyclonal at the outset (Figure 3C-D, Supplementary Table 5). In transplant mice M2, M3, M4 respectively, the dominant clone in the metastatic site arose from a minor primary clone (Figure 3C-D). Specifically, clone B, comprising 96.2% of the axillary metastatic site, formed only 1.8% of cells in the corresponding primary tumour of mouse M3. Similarly, clone P, comprising 20.8% the right axillary metastatic site, made up a tiny proportion of cells (1.8%) in the corresponding primary tumour of mouse M2. And clone K from mouse M4 reached 46% in the metastasis, and formed 0.9% in primary tumour. Several metastatic sites exhibited similar internal clonal proportionality, for example, clone D (D: 23% in primary, and E subclone of D: 40.8% in metastasis) in mouse M1 (Supplementary Table 5). Across all replicates, the clonal fraction of a primary clone appeared uncorrelated with its representation in metastases (Figure 3C-D). Taken together, these data suggest that metastatic outgrowth in Pt2-SA535 may arise from rare primary CNA clones, uncoupling metastatic fitness from primary-site dominance, reflecting a fitness cost at primary implant sites.

**Figure 3:**
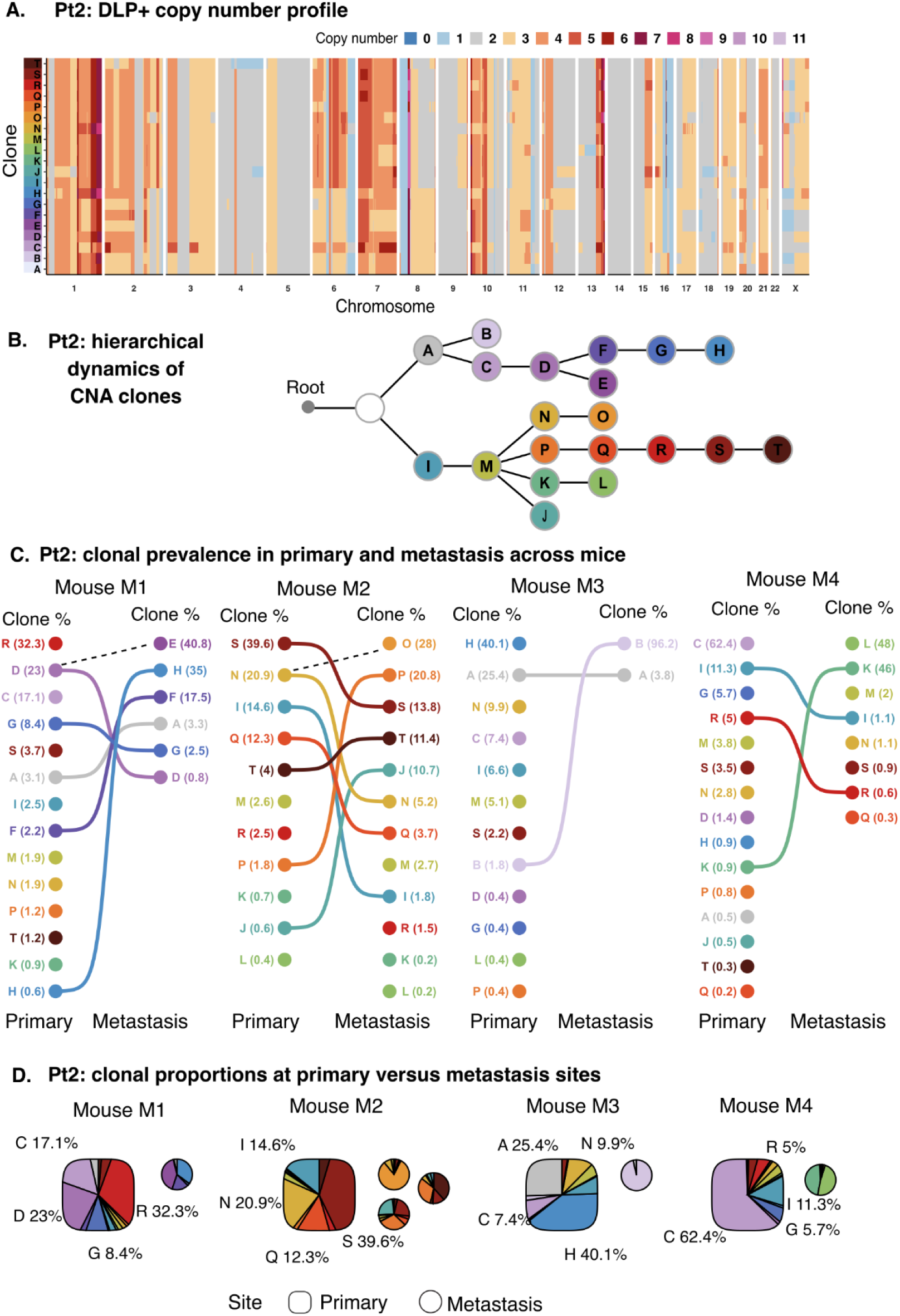
Pt2-SA535 clonal composition of transplant replicates. Pt2 exhibits polyclonality and unstable copy number alteration events. A) Heatmap of median copy number profile of each clone across chromosomes from the results of copy number analysis for Pt2. Colours denote copy number values. B) Hierarchical map of clones generated from the phylogenetic tree, output of copy number analysis. Each node is a clone. C) Bipartite graph showing the mapping between clones in primary (left) and metastasis (right) sites of 4 different mice. Colours denote clonal identities. Values denote clonal proportions. Only links between clones with at least 5% of cell representation in primary or metastasis samples are shown. Dash lines: link between sub-clones that exist only in the metastasis sample with parent clones in the primary sample. D) Clonal compositions at primary and metastasis sites of different mice (M1, M2, M3, M4) from passage 4 (X4). Colours denote clonal identities. Primary (squircle) and metastasis (circle) sites of each mouse are shown. The list of metastasis sites and clonal proportions is noted in Supplementary Table 5.

We next characterized Pt1-SA919, a TNBC with relatively low genome instability^2^, reminiscent of CNA-quiet IntClust4 ER-subtype of TNBC from the previous studies^3,29^. For Pt1-SA919, we analyzed 7 primary tumours and 7 metastasis samples from passages X3, X4, and X7 using scWGS DLP+, yielding a total of 7,074 high-quality single-cell genomes, with a median and standard deviation of 482 ± 312 single cells per sample (Supplementary Table 4). High-quality cells were used to construct copy number-based clonal structures from major clades and phylogenetic trees (see Methods) (Figure 4 and Supplementary Figure 2). Across all passages and sites, Pt1-SA919 displayed a simple clonal structure with three major CNA-defined clones (A, B, and C, Figure 4A). Clones differed from each other by CNA mutations on chromosomes 7, 5p and 10p. Clone A had a diploid chromosome 7, while clones B and C exhibited gains of whole chromosome 7 (median CN 3.0, gain in 99.6% of high-quality mapped genomic regions between clone C vs. clone A, 120 MB length, and gain in 98.8% between clone C vs. clone A, 119 MB length). Clone C contained additional CNA amplifications on chromosomes 5p (median CN 4.0, 25.8% of genomic regions with gain event, 35 MB length) and 10p (median CN 4.0, 30.3% of genomic regions with gain event, 31.5 MB length) (Figure 4A). Moreover, we extracted allele-specific copy number profiles (ASCN) generated with the SIGNALS package^2^, which showed concordance between copy number data and allele-specific copy number data, confirming the existence of only three major clones, A, B, and C, in Pt1 (Supplementary Figure 3). The clonal composition of the Pt1-SA919 tumour evolved markedly across passages X3, X4 and X7, with the earliest passage X3 comprising the A:100%, X4 comprising the A:98.8%, and B:1.2% - a minor proportion of B, respectively, with C undetectable. By passage X7, clone C contained a small proportion at primary sites and was the major clone in several metastatic tumours. Clone B, expanded from 0.2% at passage X4 to 98.3% at passage X7. Clone A was dominant at early passages but declined in proportion from 99.8% at passage X4 to 1.1% at late passage X7 (Figure 4B).

**Figure 4:**
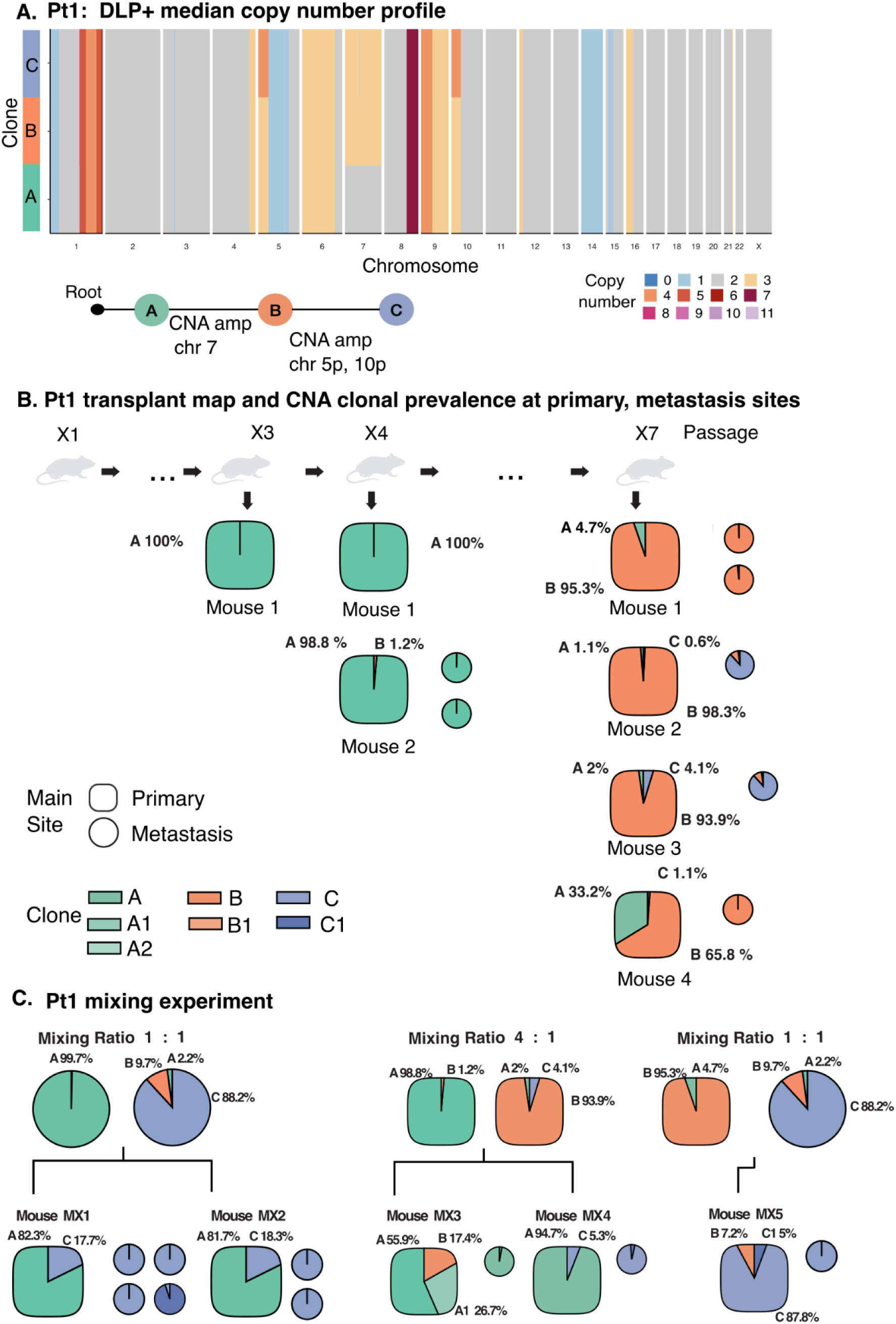
Single-cell genome sequencing DLP+ results reveal mono-clonal fitness and dominance marked by clonal sweeps in Pt1-SA919. A) Copy number analysis for Pt1-SA919. Heatmap of median copy number profile of each clone across chromosomes. Colours denote copy number values. Copy number mutational events between clone A (green) vs. clone B (orange) and C (blue) are in chromosome 7, whereas those between A and B vs. C are in chromosomes 5 and 10. B) Clonal compositions at primary and metastasis sites of different mice from passages X3, X4, and X7. Colours denote clonal identities. Primary (squircle) and metastasis (circle) sites of each mouse are shown. C) Mixing different combinations of Pt1’s primary and metastatic tumours with different clonal compositions to evaluate clonal fitness.

To evaluate how the metastatic potential of individual clones over time in Pt1-SA919, we determined the clonal cell fractions at the primary implant and metastatic sites for all primary passages X3, X4, and X7 (Supplementary Table 5), from 11 primary implants and 6 metastases. In the X3 primary passage, clone A dominated at the primary site (100% A and no presence of B, C) and no metastases were detected in any of the 3 primary site replicate transplants (0% metastatic reproducibility) up to 1 year after transplant. In primary passage X4 (A:99.1%, B:0.9%, C:0%), metastases developed in 50% of transplants (2 of 4 replicate transplants) (Supplementary Table 1). With clone A dominating both primary and metastasis sites (Figure 4C), suggesting that between X3 and X4 passages, clone A gained metastatic competence via CNA-independent processes. By passage X7, the metastatic phenotype was fully penetrant (4/4 primary transplants developed distant metastases, Supplementary Table1). At this time (passage X7), primary tumours were dominated by clone B (mean +/- sd proportions of B is 88.3 +/- 15.1% across all 4 primary sites), (Figure 4C), in addition to minor representation of clone A (10.2 +/- 15.4%), (Supplementary Table 5) and clone C ( 1.9 +/- 1.9%, detected only in 3 of 4 primary transplants at X7). In two primary transplants with only minor proportions of clone A (1.1% and 33.2%), the respective metastases were composed entirely of clone B. Strikingly, in two primary transplants (mice M2, M3) with minor proportions of clone C (C:0.6%, C:4.1%), metastases were dominated by clone C (88.2%, and 96.3% respectively) with only a minor contribution of clone B (9.7%, and 3.4%) and trace proportions of clone A (2.2%, and 0.3%, respectively) at supra-spinal and ventral-spinal metastatic sites (Figure 4C). Similar to Pt2-SA535, the emergence of clonal dominance from a minor clone suggested that clone C had developed additional metastatic potential through clonal evolution, establishing a CNA-based route to metastasis in Pt1-SA919.

First experiment: mixing ratio 1:1. Clonal prevalence for primary tumour and metastatic tumour revealed from copy number analysis. Clone A dominant metastatic samples were mixed with clone C dominant metastatic samples in equal proportion. Colours denote clonal identities. Primary (squircle) and metastasis (circle) sites of each mouse are shown.

Second experiment: mixing ratio 4:1. Clone A dominant primary samples were mixed with clone B dominant primary samples in a 4:1 ratio. Colours and shapes are denoted as (A).

Third experiment: mixing ratio 1:1. Clone B dominant primary sample was mixed with clone C dominant metastatic sample in equal proportion. Colours and shapes are denoted as (A). Main clones: A, B, C. And subclones A1, A2, B1, C1 are clones with the major events as A, B, C, but contain subtle additional events at several chromosome regions.

### Competitive retransplantation identifies copy number clone-specific metastatic potential

Prompted by the observations above, we set out to test the association between clonal fraction and metastatic reproducibility. We conducted mixture experiments in which clonal fractions were reset by either equal-proportional or strongly disproportional mixtures that disfavored metastatic clones, followed by retransplantation at the orthotopic site and single-cell sequencing of the emergent primary and metastatic tumours. Since the Pt1-SA919-X7 passage suggested apparent clonal dominance and a preference for metastasis by clone C, we first combined the dominant clone C metastasis sample with the clone A dominant metastatic sample in an equal cell ratio (mouse MX1 and MX2, Figure 4C, Supplementary Table 6). Strikingly, in both these cases, clone A re-emerged as dominant in the primary sites (A:82.3% and A:81.7%), with clone C as a minor proportion (C:17.7% and C:18.3%). In contrast, in both cases, the minor primary site clone C again dominated the metastases (C:100% in 3 metastasis sites, and C:5%, C1 subclone of C: 95% of mouse MX1, and C:100% in 2 metastasis sites from mouse MX2) (Supplementary Table 6). This shows that clone C, while having metastatic potential, is negatively fit at the primary site. To further test the dominance in metastatic potential, we conducted a second mixture transplant in which a 4:1 mixture of clone A dominant (A:98.8%, B:1.2%, C:0%) and clone B dominant cells (A:0%, B:93.9%, C:6.1%) were mixed (Figure 4C, Supplementary Table 6). In the latter clone B donor population, clone C comprised a tiny amount of cells and was thus diluted in the primary site transplant mixture to less than 1% cells (mouse MX3, Figure 4C). In two transplants, one primary site exhibited only clone A and B cells, with no clone C detected, and the corresponding metastasis was a clone A:B mixture(mouse MX3, A:55.9%, A1-subclone of A: 26.7%, B:17.4%, C:0%). In the second transplant, at the primary site, the majority of cells were clone A, and a minor population was clone C (mouse MX4: A: 94.7%, B: 0%, C: 5.3%). As predicted, the corresponding metastasis site comprised 96.4% clone C. This strongly emphasizes the dominance of the clone C metastatic phenotype. Finally, to test the dominance of clones B and C, we conducted a further ∼1:1 mixture transplant of clone B and clone C (Figure 4C, Supplementary Table 6). The primary site tumour arising comprised 87.8% clone C, and the corresponding metastasis was 100% clone C (mouse MX5, Figure 4C). Taken together, the data point to a fitness inversion of CNA-defined clones in this transplant setting, where clone A was fitter in the orthotopic primary site compared with clone C, whereas clone C exhibited much stronger metastatic potential but weaker fitness at the primary site.

### Transcriptional analysis of model PDX metastatic system reveals clonal identity-dependent transcriptional modulation and metastasis driver genes

Since Pt1 clones (A, B, C) revealed an inverse relationship between primary site fitness and metastatic potential as copy gains increased going from clone A to C (Figure 4A&C), we used to RNA-seq to investigate how gene expression changes in a copy number-dependent manner in clones A, B and C for their metastatic vs primary site transcriptional reprogramming (Figure 5, Supplementary Figure 4,5, Supplementary Table 7,8). After sequencing bulk RNA-seq of quasi-monoclonal samples of clones A, B and C, we filtered and normalized the gene expression across samples (see Methods). Gene dosage changes via somatic variations (SVs) can drive potential transcriptional changes across tens or hundreds of genes, depending on the SV size, in a tandem manner on individual chromosomes, referred to herein as “in-cis”, referring to copy number genotype-dependent in-cis genes. The differential expression (DE) analysis contrasting clone C vs B in-cis regions revealed the majority of genes, 90%, displaying up-regulation in gene expression, as expected due to copy-gain observed for genes on chromosomes 5 & 10 in clone C in comparison to clone B (Figure 5A). Similarly, comparison of clones B vs A and C vs A revealed that the majority of the ‘in-cis’ genes, 71%, and 57% respectively, displayed a positive tendency of up-regulation in gene expression, and gain in copy number values. Moreover, instead of relying on DE genes, we also used normalized expression values for all the DE genes with copy gains as observed on chromosomes 5, 7 and 10 that define the genotypic differences amongst clones A, B, and C. The evaluation of absolute normalized expression over the single copy gain regions revealed modest (Figure 5B) but significant increase (KS test, Figure 5B) in expression over the majority, i.e. 4/6 individual comparisons for chromosomes 5, 7 and 10 across clones (median gene expression values in metastasis group greater than in primary group in all 6 comparisons). Chromosome 1 contains several genomic regions with copy number change (2.5 MB length) with high amplification copy number status (median CN value 10.0 in clone A, and 9.0 in clone B, C) and low read mapping, thus making it difficult to quantify clonal difference in copy number with confidence; hence, this chromosome was excluded from DLP+ clonal analysis. As expected, there is no significant change in transcript level due to the change in the copy number of these regions of this chromosome. Hence, overall, the magnitude of gene expression positively correlates with copy number changes across clones. Moreover, these results highlight the significant downstream impact of SVs on the transcriptional landscape of clones, which can drive metastatic versus primary site fitness.

**Figure 5:**
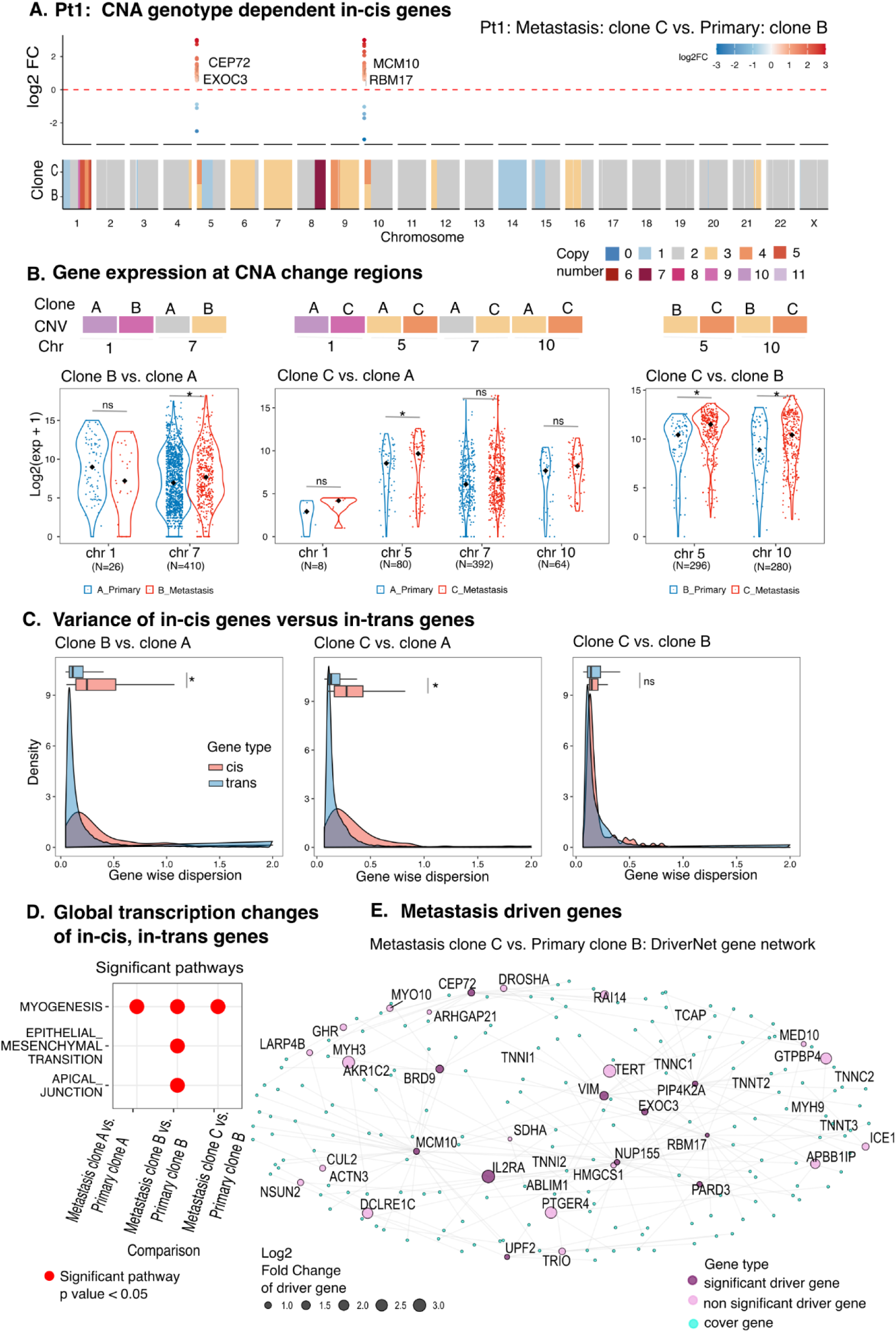
Transcription changes of PDX model Pt1-SA919 are positively correlated with copy number variations. A) Track plot showing an example of CNA genotype dependent in-cis genes. Differentially expressed in-cis genes with selected conditions: abs(log2FC) > 1, p-value < 0.05 (top). Colour gradient represents the magnitude of the log2 fold change. Median copy number profiles of clone B and clone C across chromosomes from copy number analysis (bottom). Colours denote copy number. Examples of in-cis genes are labelled, i.e, the CEP72 gene with up-regulated gene expression and gain in copy number values from 3 to 4 at chromosome 5 of its overlapping genomic regions with clonal genomic regions. B) Gene expression at chromosome positions with copy number changes and regulated in-cis genes. Median copy number values in genomic regions of clones from DLP+ data (top) and log2(normalized expression + 1) of bulk RNA-seq gene expression values (bottom) are shown. Each blue, red dot depicts one gene. The black dot indicates the median value of each group. Primary and metastatic conditions are shown in blue and red, respectively. Genes have greater gene expression values in metastasis compared to primary bulk RNA-seq gene expression and gain in copy number values of clones at their overlapping genomic regions. KS statistical test with * is significant with p value < 0.05, and ns: not significant. The number of genes taken into account for a given chromosome is noted on the x-axis. C) Distributions of gene-wise dispersion of in-cis (red) and in-trans (blue) genes from 3 differential gene expression comparisons: metastasis clone B vs. primary clone A, metastasis clone C vs. primary clone B, metastasis clone C vs. primary clone B. In-cis genes exhibit stronger dispersions/variances compared to in-trans genes. KS bootstrap statistical test with * is significant with p value < 0.05, and ns: not significant. D) Significant pathways of global transcription changes from metastasis vs. primary clonal comparisons - DE analysis. Red colour dots denote significant pathways with p-value < 0.05. E) DriverNet gene network depicting the impact of copy number changes on transcriptional networks in metastasis clone C vs. primary clone B. Significant driver genes (purple), non-significant driver genes (pink) and cover genes (cyan) that are connected/co-expressed with driver genes are depicted. The size of the driver gene circles denotes the degree of log2 fold change.

Next, we contrasted transcriptional variance of ‘CNA genotype dependent in-cis genes’ vs ‘CNA genotype independent in-trans’ differentially expressed genes to test the fixation status of copy gain genes. Overall, we expect greater variance in the in-cis group compared to in-trans genes, as in-trans genes are more plastic in nature, whereas gene dosage-based genes are fixed in the genome and require co-evolution for stabilized integration with the existing state of gene regulatory networks. We tested this hypothesis by extracting DE genes with DESeq2 and then calculating gene-wise dispersion between metastases and primary groups using a negative binomial generalized linear model implemented in edgeR (see Methods). The in-cis gene expression dispersion was significantly greater than in-trans differentially expressed genes for clone B vs A, and clone C vs A comparisons (Figure 5C, Bootstrap KS test, P< 0.05, see Methods). Very interestingly, the copy gains in clone C compared to clone B did not show significant differences in transcriptional variance for in-cis genes vs in-trans genes (Figure 5C, Bootstrap KS test, P=0.27). This lack of elevated expression divergence in the in-cis genes with respect to in-trans genes in clone C versus B suggests that the copy gain-dependent gene expression changes in clone C during its evolution from clone B are better assimilated into the existing gene regulatory networks or led to a new, stabilized state. This relatively lower divergence expression state of in-cis genes is likely to give advantage to the in-cis dependent driver genes for the fixed transcriptional fitness that can explain the strong metastatic site-specific fixed preference found for clone C as seen in the competitive re-transplantation experiments compared to other clones (Figure 4C). On the other hand, in-cis genes in clone B, during its evolution from clone A, may lack the strong fitness imparted by in-cis genes due to significantly high variance observed in in-cis genes, and thus could explain the lack of preference for metastatic vs primary site.

To identify the metastasis-specific driver genes that are dominant in each clone resulting from gene copy changes, we first identified enriched pathways by analyzing the differentially expressed genes that contrast the metastatic site landscape with the relevant primary site expression landscape. All 3 clones revealed myogenesis pathway enrichment (Figure 5D), while clone B revealed significant epithelial-mesenchymal transition (EMT) and apical junction pathways enrichment. These significant pathways associated with metastases are consistent with the findings in the literature^30,31^.

Single-copy gain is less likely to be indicated as a differentially expressed gene; however, its fixed expression state can drive downstream pathways we observe via differentially expressed genes (DEGs). Therefore, we identified statistically enriched driver genes based on the differentially expressed genes and the clone-specific copy gain information by using DriverNet, a previously described method for joint copy number and expression analysis that utilizes prior knowledge of gene co-influence graphs to assign putative driver impact scores based on DE gene node analysis. Comparing clones B vs A, C vs A, and C vs B revealed 13, 13 and 11 significant driver genes, respectively (Figure 5E, SUPP Figure 5, Supplementary Table 8). As expected, clone B vs A DriverNet analysis revealed driver genes with copy gains, which are involved in EMT fitness, such as CDH9^32^, LAMB1^33^, CD36^34^, TERT^35^, HOXA9^36^ and HOXA6^37^, and those involved in myogenesis, such as GHR^38^, CD36^39^, SEMA3A^40^, CACNA2D1^41^, GDNF^42^, GPR37^43^ and LMOD2^44^. Comparing clones C vs B revealed important invasion-associated genes such as MCM10^45^ and mesenchymal marker vimentin VIM ^46^, which may explain the strong preference of clone C for metastatic sites rather than primary sites (Figure 5E).

## DISCUSSION

The metastatic process involves multiple steps, each of which can represent an evolved phenotype of malignant cells^47^. Here, we have measured the contribution of tumour-intrinsic somatic copy number variants to metastatic fitness using a patient-derived xenograft model of spontaneous metastasis. We adopted this approach, rather than tail vein injection or other approaches that bypass metastatic steps such as microinvasion of the vasculature. The time scales of tumour growth in these treatment-naive patient lines, spanning months, rather than weeks, are thus more consistent with the long time frames of breast cancer evolution. Moreover, most of the lines studied were treatment naive, and none were derived from already metastatic cancer samples. This contrasts with cell lines that arise from extensively pre-treated, already metastatic cancers that have been extensively cultured in vitro. We also note that we undertook clonal analysis with scaled single-cell whole-genome sequencing, rather than exogenous barcoding, which admits direct assessment of the contribution of somatic CNA mutations. Gene dosage mutations are the most pervasive and dominant class of somatic variants affecting breast cancer transcriptomes^48^. The anatomical and temporal patterns of spontaneous metastasis were patient-specific, as has been indicated in earlier studies^13,17^, consistent with a strong intrinsic tumour cell determination of behaviour. Large-scale genome analysis of breast cancers has suggested that metastatic phenotypes may have a genomic underpinning from CNAs^6,7^, consistent with our observations.

A further principle of evolutionary processes is that fitness states are context-dependent, i.e., mutations that confer a positive or negative survival advantage in one environment may not operate in another^49^. In this respect, we noted that, in both patient lines studied, the clones giving rise to metastasis were often minor clones at the primary site. Competitive retransplantation experiments with downstream clonal sampling have previously been used to functionally validate inferred clonal fitness^20^. To our knowledge, the fitness of CNA-defined clones in human tumour metastasis has not been rigorously assessed with this approach. This dominance effect was reiterated in validation experiments in which metastatic clones of varying putative metastatic potential were mixed at different proportions and retransplanted. The observations showed that a hierarchy of metastatic fitness existed between CNA clones at the primary site for the line analyzed, establishing ongoing CNA mutations as one potential mechanism of acquisition of metastatic phenotypes. A trend towards less fitness at the primary site for highly metastatic clones. This suggests that some metastasis-promoting CNA mutations have negative fitness for primary tumour growth. This is analogous to observations that chemotherapy drug resistance can have a clonal fitness cost^20^.

Our study reinforces that at the molecular level, the cells with metastatic potential have EMT pathway enrichment^31^ through genetic, non-genetic processes, or a combination of both. Thus, convergence, either by transcriptional memory and/or by gene dosage mutations that pattern transcription, leads to convergence on molecular states that promote metastatic niche survival. Both processes, transcriptional landscape convergence and genome evolution, must be measured to fully understand the origins and persistence of the states conferring fitness.

We recognize that the spontaneous metastatic model transplant system is not high-throughput, and this is especially true for breast cancers that grow more slowly than some other epithelial tumours. This practically limits the understanding of patient-to-patient variation in metastatic phenotypes. Studying even a few patient lines takes many months or years, although the ability to perform perturbative and competitive repopulation assays brings quantitative validation potential. Additionally, these models, by design, lack cellular immunity and thus cannot inform on the role of immune pruning of metastatic clones and the interaction with gene dosage mutations, which is known to be relevant to cancer progression. Nevertheless, the ability to accurately measure clone fitness by retransplantation provides quantitative insights that cannot be easily obtained from single-patient samples. Understanding the general impact and the fitness costs associated with gene dosage mutations promoting metastasis will thus require future work.

## METHODS

### Experimental Procedures

#### Experimental study designs, primary and metastatic tumour detection, and serial passaging of patient-derived xenografts

Female NOD/SCID interleukin-2 receptor gamma null (NSG) and NOD Rag-1 null interleukin-2 receptor gamma null (NRG) mice were bred and housed at the Animal Resource Centre at the British Columbia Cancer Research Centre. Surgery was carried out on mice between the ages of 8 and 14 weeks. All experimental procedures were approved by the University of British Columbia Animal Care Committee (A24-0020) and Ethical Review Committee (H16-01625 and H20-00170).

#### Mammary fat pad transplant

Mice were anesthetized using isoflurane. Up to 60 μL of Lidocaine (0.25%) was injected subcutaneously for local anesthesia. Ophthalmic ointment was applied to the eyes to prevent them from drying out. The analgesic Meloxicam (Boehringer Ingelheim, DIN 02240463) was administered subcutaneously at 5mg/ml before surgery, and 0.9% sterile warm saline was administered to prevent dehydration. The hair on the flank was shaved, and a sterile skin preparation was made by Hibitane soap, followed by 70% isopropanol. A skin incision of 2-3 mm was made on the left flank of the mouse over the mammary fat pad, in a cranio-caudal direction. The mammary fat pad was exposed with forceps, and tumour cells suspended in 60 μL of a 50:50 v/v mixture of cold Matrigel:DMEM/F12 before they were injected into the mammary fat pad using a 16-gauge syringe. Skin incision was closed using Vicryl 5-0 suture with subcutaneous and simple interrupted sutures. Tumour cells were transplanted into four mice per PDX line, unless otherwise specified.

#### Surgical removal of the primary tumour

Mice were anesthetized and prepared for surgery as described above. Skin incision approximately 1-1.5cm was made over the tumour, depending on the tumour size, in the cranio-caudal direction. The tumour was dissected from surrounding tissue using Metzenbaum scissors to minimize tissue damage. The mammary fat pad attached to the tumour was removed along with the tumour. Overlying skin was removed with the tumour when the tumour was firmly attached to the skin. Skin incision was closed using Vicryl 5-0 suture with subcutaneous and simple interrupted sutures.

Selected tumours were serially passaged as previously described^28^. Briefly, tumours removed during survival surgery were finely minced with scalpels, then mechanically disaggregated for 1 minute using a Stomacher 80 Biomaster (Seward Limited, Worthing, UK) in 1 to 2 ml cold DMEM-F12 medium. Serial passaging was performed using aliquots directly from processed tumour cells or from cryopreserved materials from previous passages.

#### Organ harvesting

All mice were humanely euthanized when there was a recurrent tumour, palpable metastatic mass, symptoms suggestive of metastasis or evidence of metastasis from imaging study. Palpable tumours beneath the skin were removed first. Then a long, vertical midline incision was made on the abdomen with scissors. Any suspicious organs or masses were removed and transferred to a tube with medium (DMEM-F12) and kept in ice. The thoracic cage was opened by removing the rib cage. The heart and lungs were infused with saline via the airway before being collected into a tube. Other parts of the mouse body were thoroughly evaluated for metastasis, including the axillary area, intracranial, intraperitoneal, retroperitoneal and thoracic space. Brain, lungs, and liver were routinely harvested for histopathological review for evidence of metastasis.

#### Tissue processing for PDX tumours

A small piece of tumour tissue was removed using scalpels and fixed in 10% formalin buffered saline (Fisher Scientific, Kalamazoo, MI, USA) for histological analysis. Additional small fragments from different portions of the tissue, dissociated with a scalpel, were collected together, flash-frozen in liquid nitrogen, and stored at -80℃ for nucleic acid extraction. The remaining tissue was finely minced with scalpels, then mechanically disaggregated for 1 minute using a Stomacher 80 Biomaster (Seward Limited, Worthing, UK) in 2 ml cold DMEM-F12 medium. Aliquots from the resulting cell suspension and clumps were used for xenotransplantation or cryopreserved for single-cell analysis in DMEM-F12 medium containing 40% FBS and 10% DMSO.

#### Digestion of tumour cells for single-cell sequencing

Cryopreserved mechanically disaggregated cells/organoids were thawed rapidly in a 37°C water bath, topped up to 1.5 ml with DMEM (Sigma) and centrifuged (1100 rpm, 5 minutes), discarding the supernatant to remove DMSO from freeze media. 0.5 ml collagenase/hyaluronidase (StemCell) was added to the tissue and topped up to 1.5 ml with DMEM, then pipetted up and down to dislodge tissue pellets. The tissue was incubated at 37°C for two hours, pipetting up and down the sample for 1 minute every 30 minutes during the first hour, and every 15-20 minutes for the second hour, before centrifuging (1100 rpm, 5 minutes) and removing the supernatant. The tissue pellet was resuspended in 0.5ml trypsin, pipetted up and down for 1 minute, topped up to 1.5ml with FBS, and centrifuged (1100 rpm, 5 minutes), discarding the supernatant. 1ml dispase (StemCell) was added to the tissue pellet and pipetted up and down for 1 minute, and centrifuged for 5 minutes at 1050-1100 rpm, discarding the supernatant. Digested cells were resuspended in PBS containing 0.04% BSA to a final concentration of 1 million cells/ml. Cells were passed twice through a 70 μm filter to remove remaining undigested tissue, and this single-cell suspension was used for single-cell sequencing, DLP+.

#### Detection of metastasis

Metastasis was detected either by palpation, PET/CT or gross inspection after euthanasia and confirmed by histologic evaluation. Mice were palpated weekly until the primary tumour was detected, and then twice a week thereafter for measurement. Tumour size was measured by two technicians to reduce measurement variability. Tumours were measured in two dimensions using a digital calliper and expressed as tumour volume in mm³; defined as: [volume = 0.52 x (Length) x (Width)]. After removal of the primary tumour, mice were monitored for the development of metastasis. When mice developed palpable metastatic lesions, measurement was done twice a week until euthanasia.

#### Histopathological evaluation

Tumours and organs from mice were processed into formalin-fixed paraffin-embedded blocks. All samples were stained with hematoxylin and eosin (H&E), and slides were evaluated for the presence of tumour cells by a pathologist (Dr. Takako Kono). Tissue microarrays were constructed using duplicate 0.6mm cores extracted from formalin-fixed paraffin-embedded blocks. From each tissue microarray block, 4 μm sections were immunostained on a Ventana Discovery XT staining system (Ventana Medical Systems, AZ, USA). Sections were deparaffinized in xylene, dehydrated through three alcohol changes, and transferred to Ventana Wash solution. Endogenous peroxidase activity was blocked in 3% hydrogen peroxide. The slides were reviewed by our pathologist, Dr. Takako Kono, and the scoring method for each antibody is described in Supplementary Table 2. The positivity of biomarkers was determined as described in the table. In general, protein expression was scored visually based on the determination of staining intensity (0 – negative, 1 – weak, 2 – moderate, 3 – strong) and percentage of cells with nuclear, cytoplasmic or membranous staining (0-100%). Biomarker information was considered uninterpretable if there were no tumour cells in the cores or the cores were missing.

#### PET/CT

PET imaging experiments were conducted using a Siemens Inveon micro-PET/CT scanner. The mice were sedated with 2% isoflurane inhalation and positioned in the scanner. A baseline CT scan was obtained for localization and attenuation correction before radiotracer injection, using 80 kV X-rays at 500 μA, three sequential bed positions with 34% overlap, and 220-degree continuous rotation. This was followed by a 10-minute static PET scan. For dynamic PET scans, a 60-minute list-mode acquisition was started at the time of intravenous injection with 4-6 MBq of 18F-FDG following a baseline CT scan. The mice were kept warm by a heating pad during acquisition. The PET images were reconstructed using the ordered subset expectation maximization and maximum a posteriori algorithm (OSEM3D/MAP), using 2 OSEM3D iterations followed by 18 MAP iterations, with a requested resolution of 1.5 mm. Inveon Research Workplace was used for image viewing and analysis.

#### DLP+ library preparation

After processing tumour cells as described above, single-cell suspensions or nuclei were isolated and spotted, and libraries were prepared through the optimized DLP+ method as described previously ^2,27^. Briefly, a single-cell suspension was loaded into a contactless piezoelectric dispenser (sciFLEXARRAYER S3, Scienion) and spotted into the open nanowell arrays (SmartChip, TakaraBio). Cell dispensing was followed by enzymatic and heat lysis, and then each well was dispensed with tagmentation mix (14.33 nL TD Buffer, 3.5 nL TDE1, and 0.16 nL 10% Tween-20 or 7.5 nL Bead-Linked Transposomes (Illumina DNA Prep), 7.5 nL tagmentation buffer 1 and 15 nL nuclease-free water) followed by incubation and neutralization. Final recovery and purification of single-cell libraries were performed after 8 or 11 PCR cycles. Cleaned-up pooled single-cell libraries were analyzed using the Agilent Bioanalyzer 2100 HS kit and quantified using the Qubit dsDNA High Sensitivity kit. Libraries were sequenced at The School of Biomedical Engineering (SBME) Sequencing Core at the University of British Columbia on the Illumina NextSeq 550 (mid- or high-output, paired-end 150-bp reads), or at the Genome Sciences Centre on Illumina HiSeq2500 (paired-end 125-bp reads), Illumina HiSeqX (paired-end 150-bp reads), or the Illumina NovaSeq6000 (paired-end 150-bp reads)

#### Bulk RNA sequencing

Total RNA was extracted using the RNeasy Mini Kit (Qiagen) according to the manufacturer’s instructions. RNA quality and integrity were assessed using the Agilent 2100 Bioanalyzer and the Agilent RNA 6000 Nano Kit. Concentration was determined using Qubit. Only samples with an RNA integrity number (RIN) of 7 or higher were used for library preparation. Strand-specific RNAseq was performed using the Strand Specific Transcriptome pipeline 4.0 (Qiagen). Libraries were pooled and sequenced at the Genome Sciences Centre using an Illumina NovaSeq 6000 or HiSeq X to generate paired-end 150 bp reads.

### Computational methods

#### Tumour growth curves comparison

Upon visual inspection, it is evident that each type of cancer exhibits a different growth rate, reflecting inherent heterogeneity among individual cancers within each type. Non-metastasized cancers appear to grow faster than their metastasized counterparts. To address this heterogeneity, we applied a linear mixed-effects model^50^ from the lme4 package 1.1.34 (https://cran.r-project.org/web/packages/lme4/lme4.pdf). We used the cubic root volume of the tumours as our response variable, with metastasis status as the fixed effect, and each individual cancer as a random effect.

#### Quantification and analysis of DLP+ single-cell sequencing data

FASTQ pre-processing, sequence alignment, quality control, copy number calling and S-phase classification and filtering were performed on all libraries as detailed in our previously published paper ^27^. Briefly, cells were assigned a quality score for data quality based on a 13 features random forest classifier fitted and applied. The quality score ranges from 0 to 1, with 1 indicating a high probability that a library is high quality. Cells or nuclei with a quality score equal to or greater than a threshold of 0.75 were considered as good quality data and are considered in downstream analysis.

Copy number alterations on a per-cell basis were determined using a hidden Markov model (HMM) approach using the HMMCopy package with parameterizations ^27^. S-phase cells were identified using an automated classifier trained using cell-cycle sorted cells. Features from HMM output were used to identify cells most likely in the early or late phase of genome replication, and these S-phase cells were removed from the analysis, as they can induce strong GC bias and interfere with downstream analysis ^27^. To further enable phylogenetic inference, 5-10% of cells with the highest average copy number state jumps were removed. Upon inspection, these cells included both early- and late-dividing cells that were not captured by the S-phase classifier. Cells with low coverage (< 50 K total mapped reads) were filtered. Mouse cells were filtered from xenograft libraries using the fastqscreen method ^51^. Briefly, single cells were aligned to human and mouse references; if a cell had higher read counts in the human alignment, it was assigned to the human cell group. In contrast, cells with higher mouse cell counts were classified as mouse cells and removed from analysis.

#### Phylogenetic tree reconstruction, tree cutting, and tree visualization

To characterize and measure cell subpopulations and infer evolutionary relationships among cells, we applied Sitka, a single-cell Bayesian tree-construction method (version 1.0) 16. The input is single-cell copy number states from good-quality cells, and the output is the HMMCopy package as described above. The Sitka method converts copy number values into copy number binary change point variables. The output of this method is a phylogenetic tree in which cells are placed at the tree leaves, with a branching structure representing their ancestry.

Tree-cutting method: from the obtained phylogenetic tree, cells are assigned to clones. A clone is a group of cells in a neighbourhood, with genomic homogeneity in copy-number values or point mutation events. To construct clones, we first build a graph from the phylogenetic tree topology, then identify cell-connected components using the Leiden clustering method from the input graph ^52^.

Tree visualization: to visualize the branching structures, position of clades in the tree, cells in the neighbourhood area from the same clone in the tree are replaced by a circle with a clonal label annotation. Clones in the adjacent branches, or sub-clones, are denoted by an angle connected between them.

#### Bipartite matching

To visualize metastasis potential, i.e., a clone that is dominant or has a minor population at the primary site and becomes dominant in metastasis sites, we draw a bipartite graph using the ggbump package version 0.1.99999. First, we computed the cell proportions of each clone in primary and metastasis samples. Each clone is denoted by a node in the graph. Then, clones are ranked by the abundance - clones with higher proportions are put on the top part of the graph. A line is linked between 2 nodes in the primary and metastasis groups if 2 nodes are from the same clone. Moreover, if a new clone in metastasis is a subclone from an existing clone in primary, we manually draw a line to connect the 2 clones/nodes.

#### Bulk RNA-seq analysis

For patients Pt1 and Pt2, we captured joint tumour populations of (i) DLP+ single cell genome sequencing as described above to reconstruct a phylogenetic tree and measure copy number profiles, and (ii) bulk RNA-seq transcripts to identify differentially expressed genes. First, we aligned reads to the genome using Kallisto. We applied the Kallisto package version 0.46.1 ^53^ with human genome reference version hg38 and input of paired fastq files in order to quantify the abundance of transcripts from bulk RNA-seq data.

Differential expression analysis: to estimate the differentially expressed genes across different sample conditions, we applied the DESeq2 package ^54^ version 1.48.1. DESeq2 is based on the negative binomial distribution. The input data is a matrix of raw counts of genes x samples as output of Kallisto alignment. First, we removed library size effects by using the function computeSumFactors() from the Scran package version 1.36 ^55^. Then the obtained size factors are put back into the DESeq2 function and used as DESeq2 size factors to process the next step of estimating the dispersions and fitting a generalized linear model. The conditions are between the samples from the same clone in primary and metastasis sites, or between two clones. The output of DESeq2 is a list of genes, with log 2 fold change and p-value. Genes are considered differentially expressed genes (DE genes) if p-value < 0.05, and absolute log 2 fold change value > 1. We used the large threshold value 1 to filter genes with variance due to technical noise and get accurate DE genes. To obtain the normalized gene expression values, we divided the bulk RNA-seq gene counts by the size factors and got a log scale of expression using log2 (normalized gene expression + 1).

#### CNA genotype-dependent in-cis and independent in-trans gene detection

Genes are characterized as in-cis, or in-trans genes, based on the overlapping of their genomic regions with bin genomic regions of clonal copy number data where there are copy number changes. Gene categories are noted in our previously published paper ^21^. Briefly: (i) First, from the DLP+ copy number analysis results above, we computed the median copy number value (Cn) of copy number values across cells for each bin genomic region of each clone, using median CN values as the profile of each clone. (ii) With the list of DE genes - output from DESeq2 differential expression analysis, we used the R package annotables version 0.2.0 (https://github.com/stephenturner/annotables) to find all genomic ranges (containing start, end, width) corresponding to a given ensemble gene index. Then, the findOverlaps function from the R package iRangers ^56^ to search for any overlaps between bin genomic regions of copy number data and genomic ranges of each gene, resulting in a list of bin genomic regions and their corresponding g ene indexes at the overlapping area. (iii) A CNA genotype-dependent in-cis gene is a gene that is defined as DE from bulk RNA-seq analysis, with a change in the copy number value of pair clones at overlapping bin genomic regions. In contrast, a CNA genotype-independent in-trans gene is a DE gene without detected changes in the copy number values of overlapping genomic regions.

We further classify genes into (i) CNA correlated/ positive influence in-cis, or (ii) CNA anti-correlated/ negative influence in-cis, based on the direction of gene expression between two conditions as described in our previously published paper ^21^. CNA correlated in-cis genes included Gain-Up: gain in median copy number values between two clones in DLP+ results, and up-regulated with a positive log2 fold change value of gene expression in DE analysis between two inferred clones in bulk RNA-seq. (ii) Loss-Down: loss in copy number values between two clones in DLP+ results and down-regulated in gene expression with a negative log2 fold change value of gene expression in DE analysis. CNA anti-correlated in-cis genes include Gain-Down: gain in median copy number values and down-regulated in gene expression (iv) Loss-Up: loss, decrease in copy number values and up-regulated in gene expression.

#### Track plots

The upper panel displays a scatter plot (ggplot2 version 3.3.3^57^), with one point per DE gene. The y-axis denotes the log2 fold change value of each gene, while the x-axis represents the genomic position of each gene, calculated by extracting start, end, and chromosome genomic position of a given gene using the R package annotables version 0.1.91 (https://github.com/stephenturner/annotables). Panels are faceted by chromosomes (1–22, X). Bottom: heatmap plots of median copy number profile for a pair of clones from the output of DLP+ copy number analysis.

#### Bootstrap KS statistical test

To do statistical tests between two groups with an equal number of cells/ values, i.e Figure 5B, gene expression between metastasis and primary groups, we used the Kolmogorov-Smirnov (KS) test, the function ks.test() in R, with p-value < 0.05 means a significant difference between two groups.

For statistical tests between two groups with an unbalanced number of cells, i.e dispersion values of in-cis versus in-trans genes in Figure 5C, we used the bootstrap method. We sampled N=1000 times and, each time, randomly selected 70% of the cells/values in the small group. And selecting an equal number of cells/ values in the second, the big group. We then applied the Kolmogorov-Smirnov (KS) test (ks.test) to test the significance difference between two groups, and recorded the significant values (true or false with p-value < 0.05). The result of N sampling times and KS tests is a confidence interval ranging from 1 to 100; values < 2.75 or > 97.5 are considered outliers and significant test values.

#### Gene dispersion edgeR

To get a degree of change at the transcript level, we quantified the gene-wise dispersion. First, we input raw gene counts and library size factors to edgeR ^58^ to normalize data. And we define 2 groups: metastasis and primary samples. Then, we called the function getDispersion() from edgeR to estimate variance. From outputs, we extracted tag-wise (gene-wise) dispersion values to compare variance between groups. We selected genes from DE analysis results above to compare variations between two groups. The in-cis, in-trans gene labels are as determined as above. Then we plot the gene-wise dispersion output values from edgeR for 2 gene types: in-cis, in-trans, and visualize the data as a density plot.

#### DriverNet

Using joint tumour populations, we combined DLP+ single-cell genome sequencing and bulk RNA-seq transcript data to successfully determine CNA-dependent in-cis genes as described above. Thus, we applied the DriverNet package ^59^ version 1.48 to identify the driver genes and their mutational impacts on the gene network at the transcription level. Input data to the DriverNet package includes 3 files: (i) the first input is a genomic aberrations matrix - a binary matrix with genes in row names, and clonal labels in column names. If the genomic region of each gene overlaps with the bin genomic region of copy number data, then we assign the copy number value of the clone in this bin to the gene. After that, if there is a change in copy number value between 2 clones, the value of the gene will be TRUE for the observed clone. Here is an example: genomic region of gene G overlap with a bin genomic region of clone X with copy number value 2, we assigned the value 2 to gene G in clone X, and overlap with a bin genomic region of clone Y with copy number value 3, value 3 is assigned to gene G at clone Y. And the pair clones for observed comparison is: clone Y versus clone X. So there is a change in copy number values from 2 to 3 at gene G, then gene G is an outlier gene. We will fill in the value 1 for gene G at clone Y, and 0 for gene G at clone X to denote an outlier gene. (ii) The second input file is an influence graph. We followed the instructions in the DriverNet paper and tutorial in Bioconductor, and used the available reference gene network of functional interactions FIs ^60^ derived from Reactome, and other pathway, interaction databases, version 2022 from the Reactome website: https://reactome.org/download/tools/ReatomeFIs/FIsInGene_070323_with_annotations. txt.zip. First, we created a zero-squared matrix of observed genes by observed genes, then if there is an interaction between 2 genes in the functional interactions FIs database, we marked this interaction with the value of 1 in the matrix. We have a binary matrix of gene-gene interaction as input to the DriverNet method (iii). The third input is a gene expression matrix generated from bulk RNA-seq data with genes in row names, clonal labels in column names, and the values of TRUE and FALSE. Value TRUE if a gene is an outlier gene. Here we have a differential expression analysis (DE analysis), i.e clone Y versus clone X, and if a gene is DE gene, then we assign value TRUE to this gene at clone Y, and value FALSE to this gene at clone X. The output of DriverNet is the list of in-cis driver genes, the significant p-value, the list of cover genes - genes that connect to driver genes in the network and have strong connections, and other results values. The outputs of DriverNet are displayed in Figure 5D.

##### Pathway analysis

Pathway enrichment sets were computed from differentially expressed genes (FDR < 0.01) using the function g:GOSt() from the gprofiler2 package version 0.2.3 ^61^, the correction method is gSCS, with input values being log2 fold change values of DE genes from the results of bulk RNA-seq DE analysis, and reference sets are the hallmark gene set collection from MSigDB. Significantly enriched pathways are achieved with a p-value < 0.05.

## Author Contributions

**CRediT (Contributor Roles Taxonomy): Hoa Tran**: Methodology, Software, Validation, Formal Analysis, Investigation, Visualization, Writing, Project administration. **Gurdeep Singh**: Methodology, Software, Validation, Formal Analysis, Investigation, Writing, Visualization. **Hakwoo Lee**: Methodology, Conceptualization, Validation, Formal Analysis, Investigation, Resources, Data Curation, Writing, Visualization. **Damian Yap**: Data Curation, Investigation, Visualization, Writing - Review & Editing. **Eric Lee**: Formal Analysis, Validation, Visualization. **William Daniels:** Writing - Review & Editing. **Farhia Kabeer**: Validation, Investigation, Resources, Visualization. **Ciara H O’Flanagan:** Investigation, Resources, Validation, Writing - Review & Editing. **Vinci Au:** Methodology, Validation, Investigation, Resources. **Michael Van Vliet**: Methodology, Validation, Investigation, Resources. **Daniel Lai**: Software, Resources. **Elena Zaikova**: Software, Validation, Formal Analysis, Visualization. **Sean Beatty**: Software, Investigation, Writing - Review & Editing, Visualization, Project administration. **Esther Kong**: Data Curation. **Shuyu Fan**: Data Curation, Formal Analysis. **Jessica Chan**: Data Curation, Visualization. **Hoang Quan Dang**: Data Curation. **Teresa Ruiz de Algaza**: Validation, Data Curation, Resources. **Viviana Cerda**: Validation, Resources. **Andrew Roth**: Methodology, Supervision. **Samuel Aparicio**: Conceptualization, Methodology, Writing, Supervision, Project administration, Funding acquisition.

## ACKNOWLEDGMENTS

We are thankful to everyone in the Aparicio lab who helped with ideas, discussions and feedback on this project. We are grateful to Beixi Wang, Justina Biele, and Jazmine Brimhall for support in the single-cell sequencing wet lab experiment, as well as Jenifer Pham and Jack Yiu for their assistance in processing that data. Takako Kono provided valuable assistance with the pathological evaluation of screening TMA-stained imaging profiles. The work reported here would not be possible without the efforts of Peter Eriew in the development of the Aparicio lab’s PDX resources. We are also grateful to Sohrab Salehi, Tyler Funnell, and Marc William for valuable discussions in the analysis of the data we report here. This project was supported by the BC Cancer Foundation at BC Cancer. Samuel Aparicio holds the Nan and Lorraine Robertson Chair in Breast Cancer and is a Canada Research Chair in Molecular Oncology (950–230610 and CRC-2021-00205). This work was supported by the Office of the Assistant Secretary of Defense for Health Affairs through the Breast Cancer Research Program under Award No. ( HT9425-23-1-0820), Canadian Cancer Society Research Institute Impact program grant (705617), CIHR grants (FDN-148429, PJT-190058), Breast Cancer Research Foundation awards (BCRF-18-180, BCRF-19-180, BCRF-20-180, BCRF-21-180, BCRF-22-180, BCRF-23-180, BCRF-24-180), the Cancer Research UK Grand Challenge Program (C31893/A25050) and the Canada Foundation for Innovation (40044) to Samuel Aparicio.

## Supplementary Materials

### SUPPLEMENTARY TABLES LEGEND

**Supplementary Table 1**

List of serial passaged PDX models comprising 9 PDX lines (Pt1-9) and 5 PDX metastasis lines (Pt1-5) used in our experiment. PDX models and their respective features include: Patient ID used in this manuscript, PDX ID: corresponding SA ID and passages that used in other published papers, i.e: SA919 (X3): PDX ID is SA919, and passage number is 3 (X3). Ref: reference published papers that used transplanted tissue samples generated from these PDX lines for study. For each PDX line, we recorded the time interval required for the development of primary tumors and metastases, expressed in weeks and corresponding to euthanasia time points.. Site of metastases: anatomical site in the mouse where we detected metastatic tumours i.e: paraspinal, abdominal. Frequency: number of mice that develop metastasis relative to the total number of transplanted mice for the particular patient PDX series. Treated for breast cancer before primary patient biopsy: all collected samples from patients are treatment naive. Patient variables: Patient metastasis: presence or absence of metastases and if present, anatomical site of metastases.. Age at Diagnosis: the age at which tumours were detected. Other patient details: tumour nodes info, stage of disease, grade, lymphovascular, tumour subtype: triple negative breast cancer (TNBC).

**Supplementary Table 2**

Immunohistochemistry (IHC) scores were obtained from 9 TNBC PDX lines (Pt1-9) using tissue microarray (TMA) samples. These scores included hormone receptor status markers (ER, PR, HER2) and additional stained markers. Clinical scores are based on overall staining intensity and the proportion of stained cells (0=negative, 1=weak, 2=intermediate, 3=strong). % cells: percentage of cells positively stained at each tissue sample. Scores and percentage of stained cells are evaluated by a pathologist.

**Supplementary Table 3**

PDX tumours from patients Pt1-9 were transplanted into multiple individual mice. Throughout tumour development, all mice were closely monitored, and the length and width of any detectable tumours were measured and recorded at various time points (TumorDate_Date). Tumour volume was subsequently calculated using the formula: length multiplied by the square of width (TumorVolume_num). Tumours were permitted to grow until they reached a maximum volume of 1000 mm^3, at which point they were surgically removed (Primary Tumour Resection, PTR) and the date recorded (PTR_Date). Date_length in the table refers to the interval in days between the transplant date (not shown in the table) and the date when the tumour is measured (TumorDate_Date). Each respective mouse continued to be monitored for metastases and if metastases eventually developed in the mouse, the mouse is labelled “met” and if none were detected when the experimental end-point was reached, then it is labelled “non-met” by the researcher (expert_annotation). Each tumour was identified by its unique ID, corresponding to the patient from which it originated (e.g., Pt1, Pt2) and specific transplant (Tumor_ID).

**Supplementary Table 4**

DLP+ libraries generated for Pt1: TNBC-SA919, Pt2: TNBC-SA535 indicating passage number, number of cells (initial and after different quality control filtering conditions), median reads per cell and quality metrics.

**Supplementary Table 5**

Clones and their corresponding features indicate passage number (X3, X4, X7), main sites (Primary, Metastasis), sample_metadata: sample ID - extracted tissue id from a transplant site (i.e: SA919X3XB08939), transplant mouse id (i.e mouse M6), patient id (Pt1, Pt2), PDX series - label of PDX lines in other studies (SA919, SA535), and clonal abundance (percentage), nb_cells: number of cells in each clone for Pt1-SA919, and Pt2-SA535.

**Supplementary Table 6**

Results of mixing experiment from clonal analysis using DLP+ genome sequencing data for Pt1-SA919.

Patient_ID: Pt1, SA_ID of PDX line is SA919, transplant_mouse: mouse id that we transplanted tissues into, mouse id paper: the corresponding id of mouse that used in manuscript figure 5 to make figure simple, main site: sites of development in the transplant mouse: primary or metastasis, origin: primary or different metastasis transplant sites such as axillary, ventral spinal, supra spinal, etc. tumor_ID: id of tumour in database, detail_mixing_experiment: detail of mixing experiment i.e: Metastasis X4 (X0847-2164 Axim) + Metastasis X7 (X0847-2112252 spinal met), 1:1 means selecting tissue from metastasis site axillary of main experiment, from mouse X0847-2164 and mixed with tissue from metastasis site spinal of main experiment, from mouse X0847-2112252 at passage X7, with ratio 1:1, the transplant tissue samples. expected: the fraction of cells at each clone that we expect as result of mixing experiment at transplant sites, mixing_ratio: the mixing ratio of cells from tissue before transplanting into mouse, sample_id: sample id of harvested tissue sample in database, nb_filtered_cells: number of good quality cells after quality control from DLP+ sequencing results, nb_total_cells_raw: number of total cells achieved from DLP+ sequencing results, prop_cell_clones: proportion/ percentage of cells in each clone of mixing experiment results, main_clone: the dominance clone in each sample - output of copy number clonal analysis for these samples, jira_ticket, library_id, AT_ID: ids corresponding to these samples in database.

**Supplementary Table 7**

List of achieved genes results of bulk RNA-seq DE comparison between metastatic clone versus primary clone, for Pt1 (see Figure 6) including genes name: ensembl gene id, gene symbol; and statistical outputs of DE analysis from DESeq2: log2 fold change (logFC), PValue: p-value, and the DE comparison, i.e: Pt1 X4 Met:A vs. Pri:B: means tissue samples from PDX lines patient Pt1, passage X4, between metastasis cells in assigned clone A sample versus primary cells in assigned clone B sample.

**Supplementary Table 8**

List of driver genes and details of analysis from applying DriverNet method on DE genes from the result of bulk RNA-seq DE analyses. DE_comparison: the DE analysis comparison, i.e: Bmet_Apri: samples from assigned clone B metastasis sites versus samples from assigned clone A primary sites. Gene_type: in-cis, we focus on the copy number dependent in-cis genes. Rank: rank gene based on influence degree of this driver gene to other genes in the network. P-value: p value of statistical test based on bootstrap testing. Cover_genes: the list of genes in gene networks that have direct connection to given driver genes. Median_copy_number values for clones: the value of median copy number at overlapping region of bin genomic region at given clone from copy number data. Other details for each gene: gene symbol, biological description, ens_gene_id.

### SUPPLEMENTARY FIGURE

**Supplementary Figure 1:**
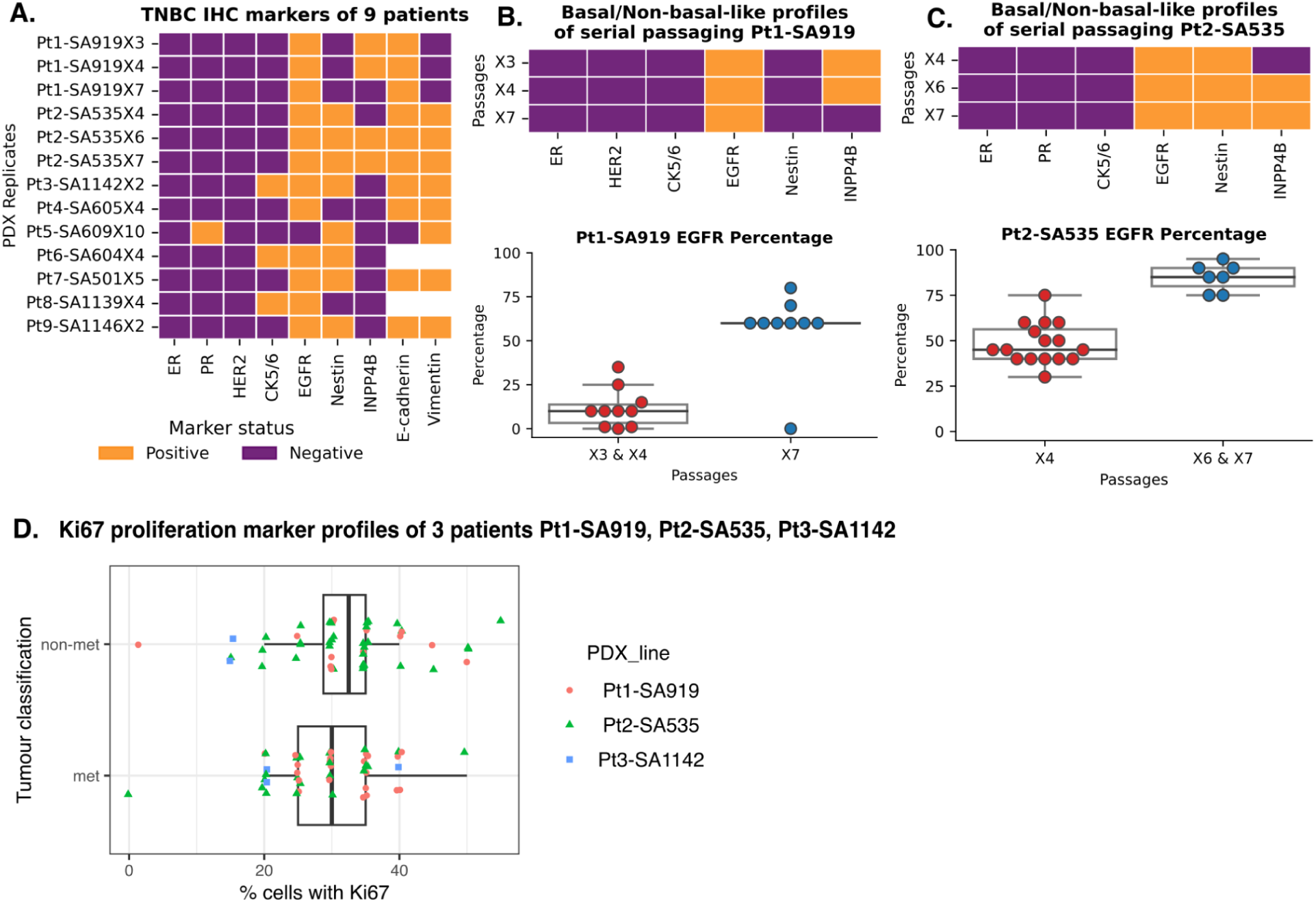
Histology, basal-like marker expressions to assess the metastasis development in multiple passages of TNBC PDX lines Pt1-9. A) Primary and metastatic tumours were assessed using immunohistochemistry (IHC) for major TNBC and cancer markers. All PDX lines are negative for ER, PR and HER2 confirming hormone receptor TNBC subtype status, except for Pt5 which showed weakly positive for PR. B) Phenotype change from non basal-like to basal-like occurred upon serial propagation of Pt1-SA919 tumours from passages X3, X4 to X7. In contrast, C) for Pt2-SA535, phenotype changes from basal-like to non basal-like from passages X4 to X6, X7. Percentage of cells positively stained with EGFR also changed upon serial passaging. D) Percentage of cells positively stained with Ki67 proliferation marker in 3 patients Pt1-3. Non met: primary transplant sites/tissue samples without metastasis development. Met: with metastasis development. PDX transplants that eventually developed metastases showed a lower fraction of proliferating cells than those that did not metastasize.

**Supplementary Figure 2:**
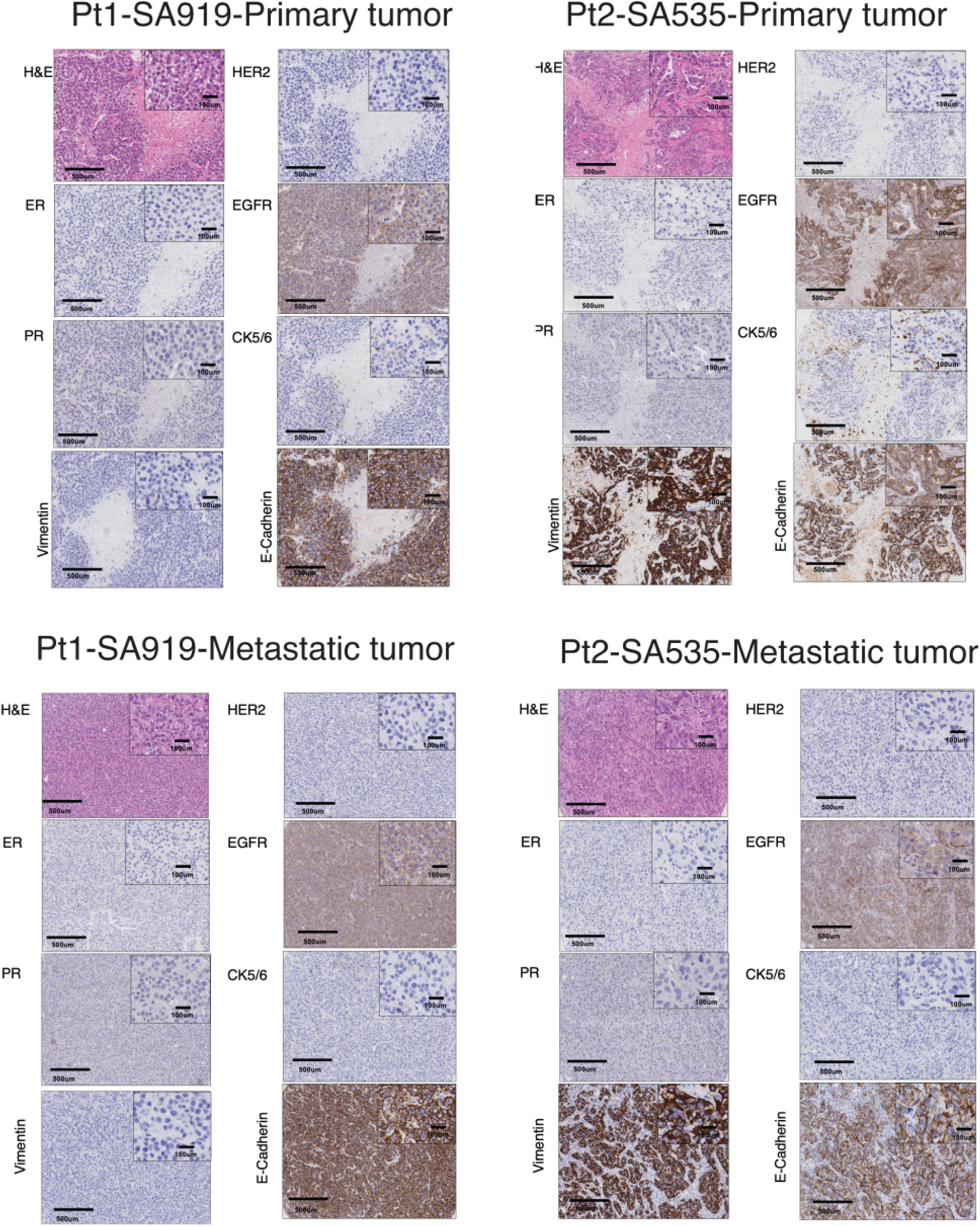
Immunohistochemistry (IHC) for TNBC and cancer markers of 5 metastatic developed PDXs. Histology images of primary and metastases tumours in Pt1-SA919 and Pt2-SA535. H&E images show primary tumours and metastasis samples (scale = 500 um for the whole sample view) with magnified view at top right in each image (scale=100 um). Notably, one representative passage and sample was picked for each PDXs here, for clinical scores details of full passages are in Supplementary Table 2.

**Supplementary Figure 3:**
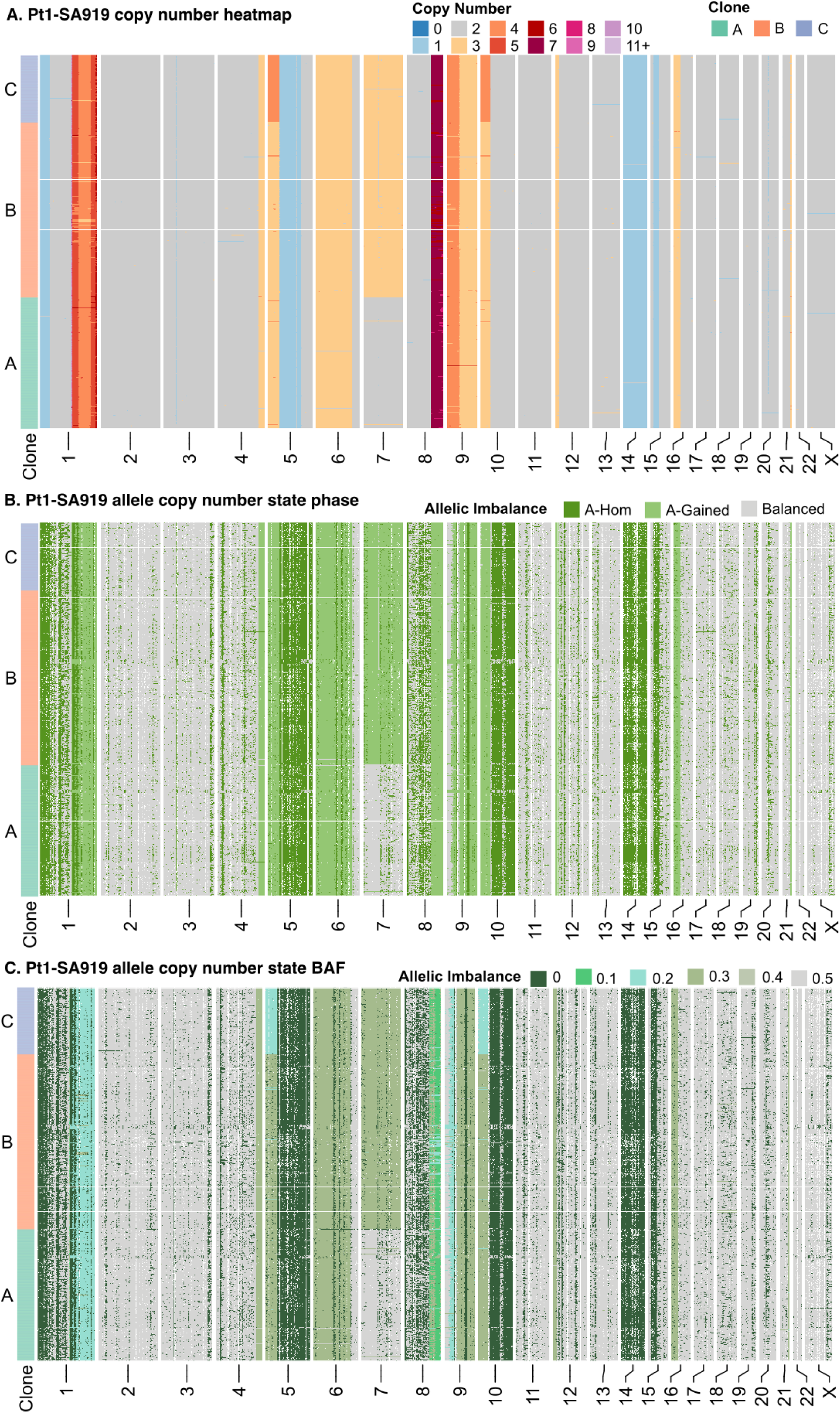
Validation of monoclonal genomic events in Pt1-SA919 using allele specific copy number data. Data are from the same list of cells (N=5471 high-quality cells common between 2 data formats copy number alteration cell profiles (CNA), and allele specific copy number cell profiles (ASCN), and 4362 genomic high quality filtered bins that share between 2 data formats) cells are at rows, and bin genomic regions at the columns. A) Copy number heatmap of Pt1 with CNA amplification events at chr 7 between clone B,C versus clone A and CNA amplification events at chr 5p, 10p between clone C versus clone B. B) Corresponding allele copy number state phases of these cells with A-gained events at chr 7 between clone B,C versus clone A. C) Corresponding allele copy number state BAF denotes the frequency of A-allele, and B-allele in each cell. Clone B, C both exhibit A-gained events but at different rates of 0.2 in clone C, and 0.3 in clone B at chr 5p, 10p. These events are in the concordance with the copy value 3 in clone B, and gain of copy number value 4 at clone C from the copy number analysis result in panel A.

**Supplementary Figure 4:**
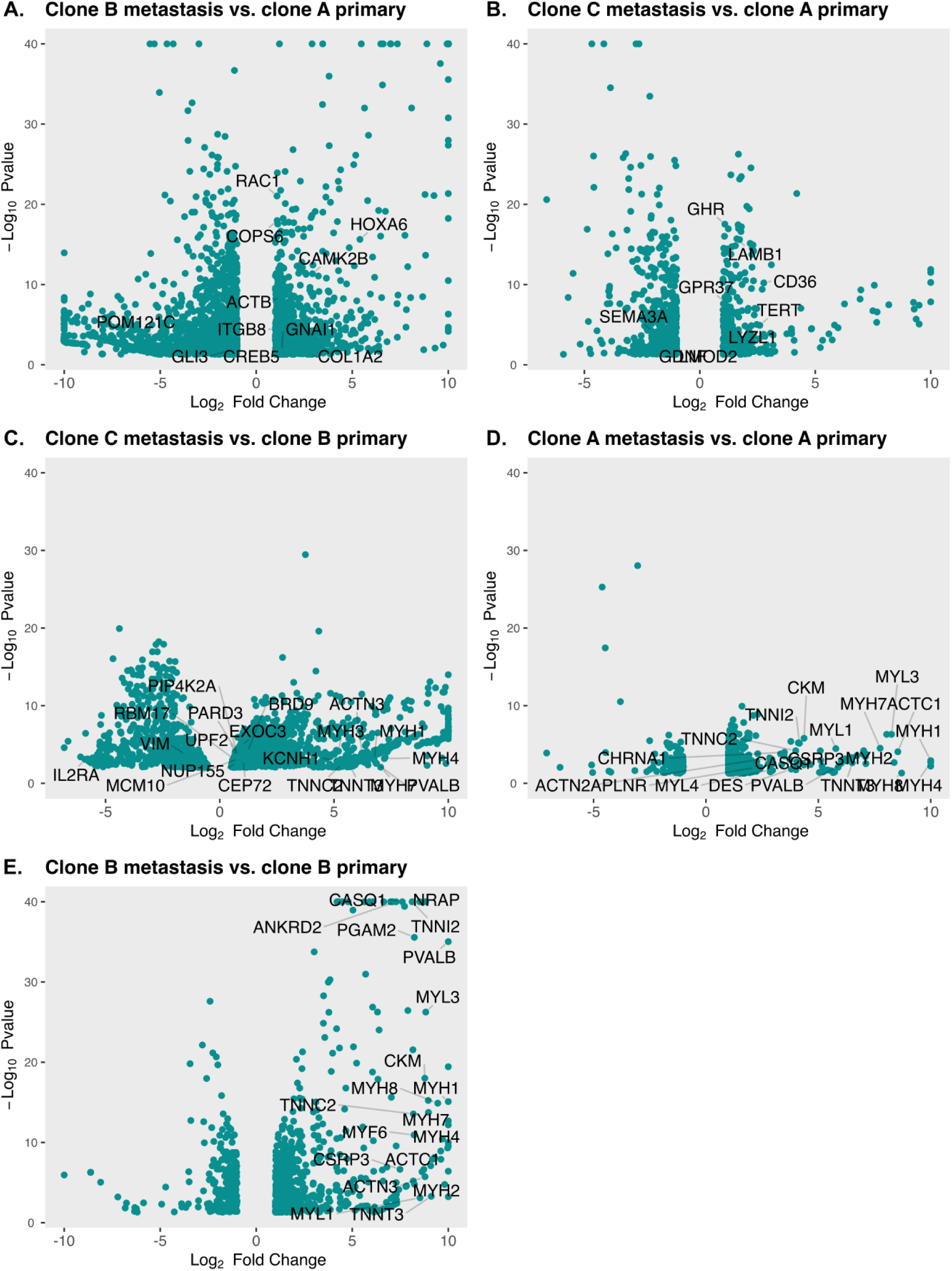
volcano plots of differential expressed analysis results applying DESeq2 method from Figure 6. Each dot is one gene, x-axis: log2 fold change results, y-axis: -log10 of P value. For comparison, clone B metastasis versus clone A primary: selecting all samples from metastasis site and assigned to clone B, and versus all samples from primary sites, assigned to clone A. In panel A, B, C genes as output of DriverNet are annotated in visualization. In panel C, D, E the top 15 genes from significant pathways are annotated.

**Supplementary Figure 5:**
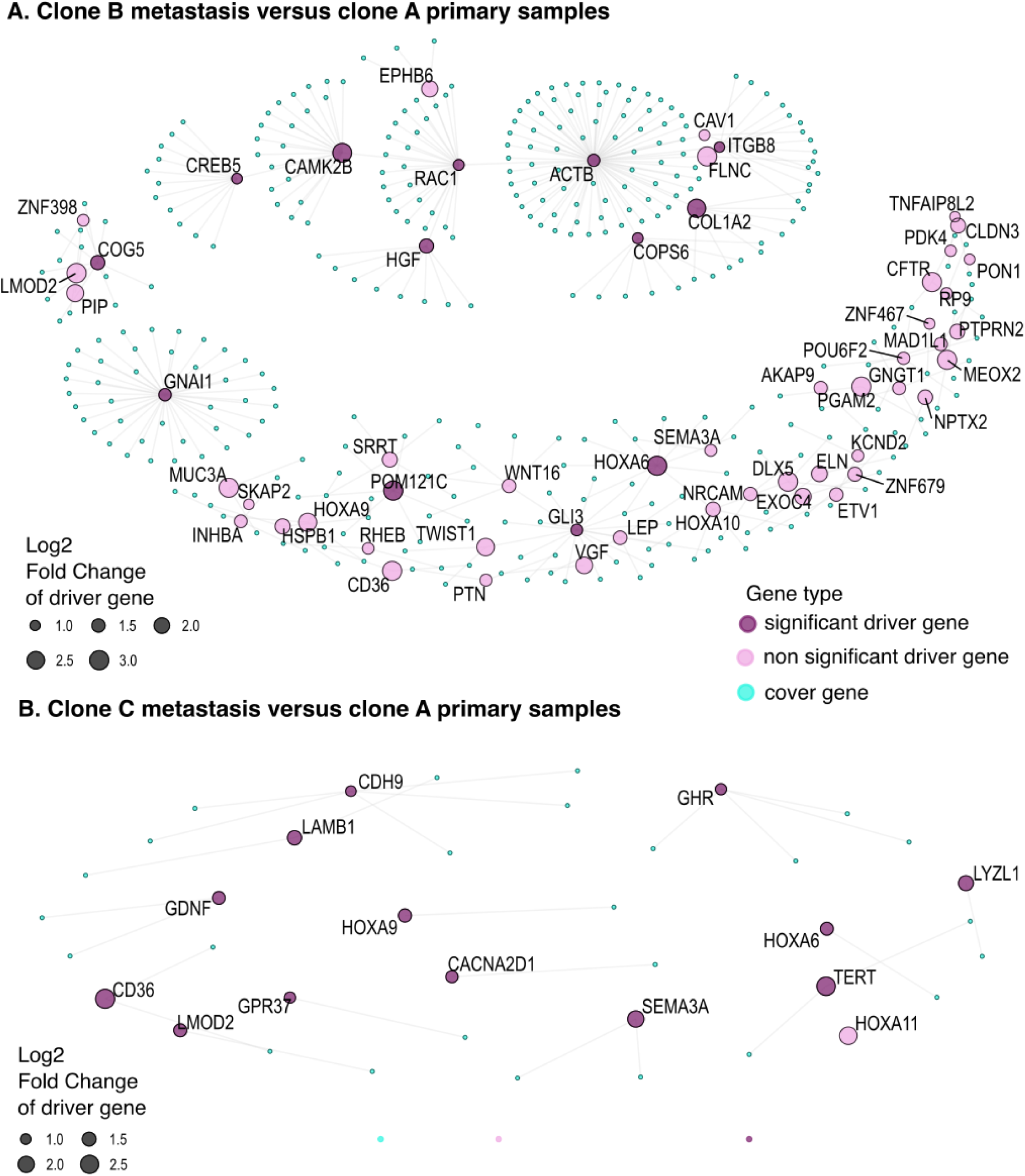
DriverNet gene network depicting the impact of copy number changes on transcriptional networks in metastasis. Significant driver genes (purple), non significant driver genes (pink) and cover genes (cyan) that are connected/co-expressed with driver genes are depicted. The size of driver genes circles denote the degree of gene expression at log2 fold change.

## Notes

### Competing Interest Statement

S. Aparicio is cofounder and shareholder of Genome Therapeutics, uncompensated advisor to Chordia Therapeutics and advisor to Sangamo Therapeutics. The other authors declare no competing interests.

## References

1. López, S., Lim, E.L., Horswell, S., Haase, K., Huebner, A., Dietzen, M., Mourikis, T.P., Watkins, T.B.K., Rowan, A., Dewhurst, S.M., et al. (2020). Interplay between whole-genome doubling and the accumulation of deleterious alterations in cancer evolution. Nat. Genet. 52, 283–293. 10.1038/s41588-020-0584-7.

2. Funnell, T., O’Flanagan, C.H., Williams, M.J., McPherson, A., McKinney, S., Kabeer, F., Lee, H., Salehi, S., Vázquez-García, I., Shi, H., et al. (2022). Single-cell genomic variation induced by mutational processes in cancer. Nature 612, 106–115. 10.1038/s41586-022-05249-0.

3. Curtis, C., Shah, S.P., Chin, S.-F., Turashvili, G., Rueda, O.M., Dunning, M.J., Speed, D., Lynch, A.G., Samarajiwa, S., Yuan, Y., et al. (2012). The genomic and transcriptomic architecture of 2,000 breast tumours reveals novel subgroups. Nature 486, 346–352. 10.1038/nature10983.

4. Rheinbay, E., Nielsen, M.M., Abascal, F., Wala, J.A., Shapira, O., Tiao, G., Hornshøj, H., Hess, J.M., Juul, R.I., Lin, Z., et al. (2020). Analyses of non-coding somatic drivers in 2,658 cancer whole genomes. Nature 578, 102–111. 10.1038/s41586-020-1965-x.

5. Stephens, P.J., Tarpey, P.S., Davies, H., Van Loo, P., Greenman, C., Wedge, D.C., Nik-Zainal, S., Martin, S., Varela, I., Bignell, G.R., et al. (2012). The landscape of cancer genes and mutational processes in breast cancer. Nature 486, 400–404. 10.1038/nature11017.

6. Rueda, O.M., Sammut, S.-J., Seoane, J.A., Chin, S.-F., Caswell-Jin, J.L., Callari, M., Batra, R., Pereira, B., Bruna, A., Ali, H.R., et al. (2019). Dynamics of breast-cancer relapse reveal late-recurring ER-positive genomic subgroups. Nature 567, 399–404. 10.1038/s41586-019-1007-8.

7. Bertucci, F., Ng, C.K.Y., Patsouris, A., Droin, N., Piscuoglio, S., Carbuccia, N., Soria, J.C., Dien, A.T., Adnani, Y., Kamal, M., et al. (2019). Genomic characterization of metastatic breast cancers. Nature 569, 560–564. 10.1038/s41586-019-1056-z.

8. Weigelt, B., Peterse, J.L., and van ’t Veer, L.J. (2005). Breast cancer metastasis: markers and models. Nat. Rev. Cancer 5, 591–602. 10.1038/nrc1670.

9. Anderson, R.L., Balasas, T., Callaghan, J., Coombes, R.C., Evans, J., Hall, J.A., Kinrade, S., Jones, D., Jones, P.S., Jones, R., et al. (2019). A framework for the development of effective anti-metastatic agents. Nat. Rev. Clin. Oncol. 16, 185–204. 10.1038/s41571-018-0134-8.

10. Nguyen, D.X., Bos, P.D., and Massagué, J. (2009). Metastasis: from dissemination to organ-specific colonization. Nat. Rev. Cancer 9, 274–284. 10.1038/nrc2622.

11. Vanharanta, S., and Massagué, J. (2013). Origins of metastatic traits. Cancer Cell 24, 410–421. 10.1016/j.ccr.2013.09.007.

12. Batra, R.N., Lifshitz, A., Vidakovic, A.T., Chin, S.-F., Sati-Batra, A., Sammut, S.-J., Provenzano, E., Ali, H.R., Dariush, A., Bruna, A., et al. (2021). DNA methylation landscapes of 1538 breast cancers reveal a replication-linked clock, epigenomic instability and cis-regulation. Nat. Commun. 12, 1–13. 10.1038/s41467-021-25661-w.

13. Bruna, A., Rueda, O.M., Greenwood, W., Batra, A.S., Callari, M., Batra, R.N., Pogrebniak, K., Sandoval, J., Cassidy, J.W., Tufegdzic-Vidakovic, A., et al. (2016). A Biobank of Breast Cancer Explants with Preserved Intra-tumor Heterogeneity to Screen Anticancer Compounds. Cell 167, 260–274.e22. 10.1016/j.cell.2016.08.041.

14. Blanchard, Z., Brown, E.A., Ghazaryan, A., and Welm, A.L. (2025). PDX models for functional precision oncology and discovery science. Nat. Rev. Cancer 25, 153–166. 10.1038/s41568-024-00779-3.

15. Dobrolecki, L.E., Airhart, S.D., Alferez, D.G., Aparicio, S., Behbod, F., Bentires-Alj, M., Brisken, C., Bult, C.J., Cai, S., Clarke, R.B., et al. (2016). Patient-derived xenograft (PDX) models in basic and translational breast cancer research. Cancer Metastasis Rev. 35, 547–573. 10.1007/s10555-016-9653-x.

16. Vaklavas, C., Matsen, C.B., Chu, Z., Boucher, K.M., Scherer, S.D., Pathi, S., Beck, A., Brownson, K.E., Buys, S.S., Chittoria, N., et al. (2024). TOWARDS study: Patient-derived xenograft engraftment predicts poor survival in patients with newly diagnosed triple-negative breast cancer. JCO Precis. Oncol. 8, e2300724. 10.1200/PO.23.00724.

17. Hidalgo, M., Amant, F., Biankin, A.V., Budinská, E., Byrne, A.T., Caldas, C., Clarke, R.B., de Jong, S., Jonkers, J., Mælandsmo, G.M., et al. (2014). Patient-derived xenograft models: an emerging platform for translational cancer research. Cancer Discov. 4, 998–1013. 10.1158/2159-8290.CD-14-0001.

18. Liu, Y., Wu, W., Cai, C., Zhang, H., Shen, H., and Han, Y. (2023). Patient-derived xenograft models in cancer therapy: technologies and applications. Signal Transduction and Targeted Therapy 8, 1–24. 10.1038/s41392-023-01419-2.

19. Echeverria, G.V., Powell, E., Seth, S., Ge, Z., Carugo, A., Bristow, C., Peoples, M., Robinson, F., Qiu, H., Shao, J., et al. (2018). High-resolution clonal mapping of multi-organ metastasis in triple negative breast cancer. Nat. Commun. 9, 1–17. 10.1038/s41467-018-07406-4.

20. Salehi, S., Kabeer, F., Ceglia, N., Andronescu, M., Williams, M.J., Campbell, K.R., Masud, T., Wang, B., Biele, J., Brimhall, J., et al. (2021). Clonal fitness inferred from time-series modelling of single-cell cancer genomes. Nature 595, 585–590. 10.1038/s41586-021-03648-3.

21. Kabeer, F., Tran, H., Andronescu, M., Singh, G., Lee, H., Salehi, S., Wang, B., Biele, J., Brimhall, J., Gee, D., et al. (2024). Single-cell decoding of drug induced transcriptomic reprogramming in triple negative breast cancers. Genome Biol. 25, 191. 10.1186/s13059-024-03318-3.

22. Byrne, A.T., Alférez, D.G., Amant, F., Annibali, D., Arribas, J., Biankin, A.V., Bruna, A., Budinská, E., Caldas, C., Chang, D.K., et al. (2017). Interrogating open issues in cancer precision medicine with patient-derived xenografts. Nat. Rev. Cancer 17, 254–268. 10.1038/nrc.2016.140.

23. Aparicio, S., Hidalgo, M., and Kung, A.L. (2015). Examining the utility of patient-derived xenograft mouse models. Nat. Rev. Cancer 15, 311–316. 10.1038/nrc3944.

24. Zhang, X., Claerhout, S., Prat, A., Dobrolecki, L.E., Petrovic, I., Lai, Q., Landis, M.D., Wiechmann, L., Schiff, R., Giuliano, M., et al. (2013). A renewable tissue resource of phenotypically stable, biologically and ethnically diverse, patient-derived human breast cancer xenograft models. Cancer Res. 73, 4885–4897. 10.1158/0008-5472.CAN-12-4081.

25. Nahed, A.S., and Shaimaa, M.Y. (2016). Ki-67 as a prognostic marker according to breast cancer molecular subtype. Cancer Biol. Med. 13, 496. 10.20892/j.issn.2095-3941.2016.0066.

26. Cunningham, J.J., Brown, J.S., Vincent, T.L., and Gatenby, R.A. (2015). Divergent and convergent evolution in metastases suggest treatment strategies based on specific metastatic sites. Evol. Med. Public Health 2015, 76–87. 10.1093/emph/eov006.

27. Laks, E., McPherson, A., Zahn, H., Lai, D., Steif, A., Brimhall, J., Biele, J., Wang, B., Masud, T., Ting, J., et al. (2019). Clonal decomposition and DNA replication states defined by scaled single-cell genome sequencing. Cell 179, 1207–1221.e22. 10.1016/j.cell.2019.10.026.

28. Eirew, P., Steif, A., Khattra, J., Ha, G., Yap, D., Farahani, H., Gelmon, K., Chia, S., Mar, C., Wan, A., et al. (2015). Dynamics of genomic clones in breast cancer patient xenografts at single-cell resolution. Nature 518, 422–426.

29. Shah, S.P., Roth, A., Goya, R., Oloumi, A., Ha, G., Zhao, Y., Turashvili, G., Ding, J., Tse, K., Haffari, G., et al. (2012). The clonal and mutational evolution spectrum of primary triple-negative breast cancers. Nature 486, 395–399. 10.1038/nature10933.

30. Hogan, K.A., Cho, D.S., Arneson, P.C., Samani, A., Palines, P., Yang, Y., and Doles, J.D. (2018). Tumor-derived cytokines impair myogenesis and alter the skeletal muscle immune microenvironment. Cytokine 107, 9–17. 10.1016/j.cyto.2017.11.006.

31. Yeung, K.T., and Yang, J. (2017). Epithelial-mesenchymal transition in tumor metastasis. Mol. Oncol. 11, 28–39. 10.1002/1878-0261.12017.

32. Clay, M.R., and Halloran, M.C. (2014). Cadherin 6 promotes neural crest cell detachment via F-actin regulation and influences active Rho distribution during epithelial-to-mesenchymal transition. Development 141, 2506–2515. 10.1242/dev.105551.

33. Bera, A., and Lewis, S.M. (2020). Regulation of epithelial-to-mesenchymal transition by alternative translation initiation mechanisms and its implications for cancer metastasis. Int. J. Mol. Sci. 21, 4075. 10.3390/ijms21114075.

34. Deng, M., Cai, X., Long, L., Xie, L., Ma, H., Zhou, Y., Liu, S., and Zeng, C. (2019). CD36 promotes the epithelial-mesenchymal transition and metastasis in cervical cancer by interacting with TGF-β. J. Transl. Med. 17, 352. 10.1186/s12967-019-2098-6.

35. Wu, Y., Bian, C., Zhen, C., Liu, L., Lin, Z., Nisar, M.F., Wang, M., Bartsch, J.W., Huang, E., Ji, P., et al. (2017). Telomerase reverse transcriptase mediates EMT through NF-κB signaling in tongue squamous cell carcinoma. Oncotarget 8, 85492–85503. 10.18632/oncotarget.20888.

36. Liu, M., Xiao, Y., Tang, W., Li, J., Hong, L., Dai, W., Zhang, W., Peng, Y., Wu, X., Wang, J., et al. (2020). HOXD9 promote epithelial-mesenchymal transition and metastasis in colorectal carcinoma. Cancer Med. 9, 3932–3943. 10.1002/cam4.2967.

37. Lin, J., Zhu, H., Hong, L., Tang, W., Wang, J., Hu, H., Wu, X., Chen, Y., Liu, G., Yang, Q., et al. (2021). Coexpression of HOXA6 and PBX2 promotes metastasis in gastric cancer. Aging (Albany NY) 13, 6606–6624. 10.18632/aging.202426.

38. Sotiropoulos, A., Ohanna, M., Kedzia, C., Menon, R.K., Kopchick, J.J., Kelly, P.A., and Pende, M. (2006). Growth hormone promotes skeletal muscle cell fusion independent of insulin-like growth factor 1 up-regulation. Proc. Natl. Acad. Sci. U. S. A. 103, 7315–7320. 10.1073/pnas.0510033103.

39. Sun, J., Su, Y., Xu, Y., Qin, D., He, Q., Qiu, H., Zhuo, J., and Li, W. (2022). CD36 deficiency inhibits proliferation by cell cycle control in skeletal muscle cells. Front. Physiol. 13, 947325. 10.3389/fphys.2022.947325.

40. Tatsumi, R., Suzuki, T., Do, M.-K.Q., Ohya, Y., Anderson, J.E., Shibata, A., Kawaguchi, M., Ohya, S., Ohtsubo, H., Mizunoya, W., et al. (2017). Slow-myofiber commitment by semaphorin 3A secreted from myogenic stem cells. Stem Cells 35, 1815–1834. 10.1002/stem.2639.

41. Inoue, H., Shiozaki, A., Kosuga, T., Shimizu, H., Kudou, M., Arita, T., Konishi, H., Komatsu, S., Kuriu, Y., Kubota, T., et al. (2024). CACNA2D1 regulates the progression and influences the microenvironment of colon cancer. J. Gastroenterol. 59, 556–571. 10.1007/s00535-024-02095-x.

42. Li, L., Rozo, M., Yue, S., Zheng, X., J Tan, F., Lepper, C., and Fan, C.-M. (2019). Muscle stem cell renewal suppressed by Gas1 can be reversed by GDNF in mice. Nat. Metab. 1, 985–995. 10.1038/s42255-019-0110-3.

43. Fabre, P., Molina, T., Larose, J., Greffard, K., Généreux-Gamache, G., Deprez, A., Mokhtari, I., Pellerito, O., Duchesne, E., Dort, J., et al. (2025). Bioactive lipid mediator class switching regulates myogenic cell progression and muscle regeneration. Nat. Commun. 16, 5578. 10.1038/s41467-025-60586-8.

44. Larrinaga, T.M., Farman, G.P., Mayfield, R.M., Yuen, M., Ahrens-Nicklas, R.C., Cooper, S.T., Pappas, C.T., and Gregorio, C.C. (2024). Lmod2 is necessary for effective skeletal muscle contraction. Sci. Adv. 10, eadk1890. 10.1126/sciadv.adk1890.

45. Yang, W.-D., and Wang, L. (2019). MCM10 facilitates the invaded/migrated potentials of breast cancer cells via Wnt/β-catenin signaling and is positively interlinked with poor prognosis in breast carcinoma. J. Biochem. Mol. Toxicol. 33, e22330. 10.1002/jbt.22330.

46. Berr, A.L., Wiese, K., Dos Santos, G., Koch, C.M., Anekalla, K.R., Kidd, M., Davis, J.M., Cheng, Y., Hu, Y.-S., and Ridge, K.M. (2023). Vimentin is required for tumor progression and metastasis in a mouse model of non-small cell lung cancer. Oncogene 42, 2074–2087. 10.1038/s41388-023-02703-9.

47. Lambert, A.W., Pattabiraman, D.R., and Weinberg, R.A. (2017). Emerging biological principles of metastasis. Cell 168, 670–691. 10.1016/j.cell.2016.11.037.

48. PCAWG Transcriptome Core Group, Calabrese, C., Davidson, N.R., Demircioğlu, D., Fonseca, N.A., He, Y., Kahles, A., Lehmann, K.-V., Liu, F., Shiraishi, Y., et al. (2020). Genomic basis for RNA alterations in cancer. Nature 578, 129–136. 10.1038/s41586-020-1970-0.

49. Michor, F., Nowak, M.A., and Iwasa, Y. (2006). Stochastic dynamics of metastasis formation. J. Theor. Biol. 240, 521–530. 10.1016/j.jtbi.2005.10.021.

50. Bates, D., Mächler, M., Bolker, B., and Walker, S. (2015). Fitting linear mixed-effects models Usinglme4. J. Stat. Softw. 67. 10.18637/jss.v067.i01.

51. Wingett, S.W., and Andrews, S. (2018). FastQ Screen: A tool for multi-genome mapping and quality control. F1000Res. 7, 1338. 10.12688/f1000research.15931.1.

52. Traag, V.A., Waltman, L., and van Eck, N.J. (2019). From Louvain to Leiden: guaranteeing well-connected communities. Sci. Rep. 9, 5233. 10.1038/s41598-019-41695-z.

53. Bray, N.L., Pimentel, H., Melsted, P., and Pachter, L. (2016). Near-optimal probabilistic RNA-seq quantification. Nat. Biotechnol. 34, 525–527. 10.1038/nbt.3519.

54. Love, M.I., Huber, W., and Anders, S. (2014). Moderated estimation of fold change and dispersion for RNA-seq data with DESeq2. bioRxiv. 10.1101/002832.

55. Lun, A.T.L., McCarthy, D.J., and Marioni, J.C. (2016). A step-by-step workflow for low-level analysis of single-cell RNA-seq data with Bioconductor. F1000Res. 5, 2122. 10.12688/f1000research.9501.2.

56. Lawrence, M., Huber, W., Pagès, H., Aboyoun, P., Carlson, M., Gentleman, R., Morgan, M.T., and Carey, V.J. (2013). Software for computing and annotating genomic ranges. PLoS Comput. Biol. 9, e1003118. 10.1371/journal.pcbi.1003118.

57. Wickham, H. (2009). Ggplot2: Elegant graphics for data analysis (Springer).

58. Chen, Y., Chen, L., Lun, A.T.L., Baldoni, P.L., and Smyth, G.K. (2025). edgeR v4: powerful differential analysis of sequencing data with expanded functionality and improved support for small counts and larger datasets. Nucleic Acids Res. 53. 10.1093/nar/gkaf018.

59. Bashashati, A., Haffari, G., Ding, J., Ha, G., Lui, K., Rosner, J., Huntsman, D.G., Caldas, C., Aparicio, S.A., and Shah, S.P. (2012). DriverNet: uncovering the impact of somatic driver mutations on transcriptional networks in cancer. Genome Biol. 13, R124. 10.1186/gb-2012-13-12-r124.

60. Wu, G., Feng, X., and Stein, L. (2010). A human functional protein interaction network and its application to cancer data analysis. Genome Biol. 11, R53. 10.1186/gb-2010-11-5-r53.

61. Reimand, J., Kull, M., Peterson, H., Hansen, J., and Vilo, J. (2007). g:Profiler--a web-based toolset for functional profiling of gene lists from large-scale experiments. Nucleic Acids Res. 35, W193–W200. 10.1093/nar/gkm226.

